# Mathematical Modelling of Bacterial DNA Inversion Dynamics Uncovers an Organized Multi-Locus Response to Phage Predation in *Bacteroides fragilis*

**DOI:** 10.64898/2026.08.07.743496

**Authors:** Avigail Belansky, Naama Geva-Zatorsky

## Abstract

Phase variation enables bacteria to generate phenotypic diversity through reversible genomic DNA inversions that alter surface structures and other adaptive traits. In *Bacteroides fragilis*, multiple invertible regions regulate surface structures, including capsular polysaccharides, which shape the bacterial interactions with the host. Previous studies have shown that molecular phase-variable surface states can alter bacteriophage susceptibility in bacteria. Here, we modelled the dynamic interaction from a longitudinal gnotobiotic mouse experiment from a recent *B. fragilis* NCTC 9343–Barc2635 study. We aimed to analyze temporal patterns, region-level susceptibility, and combinatorial patterns across the 18 invertible regions, and quantify it into a dynamical framework. The model we generated revealed a structured phage-susceptibility landscape in which loci differed in effective phage-associated sensitivity and occupied distinct parameter regimes. Projecting fitted susceptibility weights onto observed “ON”-fraction trajectories showed that the temporal response was compressed into a small subset of dominant phase variable region (PVR) contributors. A two-dimensional contribution-space analysis further separated persistent contributors from rare high-impact loci, indicating that susceptibility evolves along constrained temporal paths rather than fluctuating randomly across promoter states.

Several loci contributed to the modelled susceptibility signal over time in phase-dependent patterns. Specifically, the PVR of polysaccharide F (PSF) provided a persistent contribution, with a recurring PSF-centered, phase-dependent susceptibility pattern in combinatorial scoring of pairwise, triple, and quadruple loci sets. Our results do not identify a physical Barc2635 receptor or establish direct causal infection states. Instead, they show that phage predation is associated with a structured, low-dimensional, multi-locus organization of phase variation linking region-level susceptibility, temporal contribution, and recurring promoter-state combinations.

**Highlights:** • Development of a longitudinal mathematical framework for multi-locus bacterial phase variation under phage predation.

• The mathematical modeling revealed an organized susceptibility landscape despite high-dimensional DNA inversion dynamics.

• PSF was identified as a persistent contributor to phase-dependent multi-locus combinations during phage exposure.

• Distinguished transient phase-variable responses from sustained contributors to longitudinal phage dynamics.

• Established a general framework for interpreting bacterial genomic plasticity in host– bacterium–phage systems.

## Introduction

DNA inversion-mediated phase variation is a reversible genetic mechanism that allows bacteria to diversify their gene expression by switching defined genomic segments between two orientations^1–4^ . These invertible DNA segments, also referred to as “invertons” or phase-variable regions, can reside in promoters or regulatory sequences, lie within coding regions, or span larger local genomic segments^1–8^.

Although DNA inversions are reversible, the fraction of bacterial cells carrying each orientation is not necessarily fixed at 50:50. Instead, orientation distributions can be reproducible and condition-dependent, reflecting both locus-specific switching dynamics and environmental filtering or regulation by host context, inflammation, or phage exposure^9–13^.

In invertible promoter-associated loci, one orientation can drive transcription of a downstream gene or operon and is referred to as the “ON” orientation, whereas the alternative orientation prevents, or redirects transcription and is referred to as the “OFF” orientation. In this study, we use “ON” and “OFF” according to known motifs to denote the orientation states of each invertible region, without assuming that every PVR is a validated promoter switch or that orientation directly predicts expression for all loci^5,7,10^. In this way, DNA inversions can generate reversible heterogeneity in bacterial surface architecture and regulatory state.

This mechanism is compelling in gut bacteria such as *Bacteroides fragilis*, where surface structures influence colonization, nutrient acquisition, immune recognition, and host-microbe interactions^14–17^. Genomic and molecular studies showed that *B. fragilis* contains extensive DNA inversions that regulate variable expression of capsular polysaccharides (CPS) and broader cell-surface architecture^5,6,18–20^ . The seven invertible capsular polysaccharide promoter loci, PSA, PSB, PSD, PSE, PSF, PSG, and PSH, are central examples of this architecture. Their orientations have been measured across *in vitro* growth, gut colonization, and abscess infection, and recent work has further linked CPS promoter inversion and transcriptional regulation to capsule diversity under different environmental contexts^10,21^. Together with genome-scale approaches for identifying and quantifying bacterial invertons, these studies position DNA inversion-mediated phase variation as an important mechanism by which gut bacteria modulate functionality in response to host and microbial environments^7,8^.

Bacteriophages are viruses that infect bacteria. Lytic phages replicate inside susceptible bacterial cells and lyse them and therefore phage exposure of a bacterial population can impose strong selection on bacterial surface states. This resulting phenotypic variability can be involved in viral attachment, infection, or protection^22,23^. Phage–bacteria interactions do not necessarily result in bacterial eradication, either in *vitro* or *in vivo*^24,25^. In the gut, spatial constraints, variation in bacterial susceptibility, and broader ecological heterogeneity can further promote prolonged coexistence between phages and their bacterial hosts^26–30^. Since many phages recognize bacterial surface structures, reversible changes in surface expression can alter phage susceptibility. This phenomenon was shown in several bacterial systems. For example, in culture-based experiments with *Bacteroides thetaiotaomicron*, phase-variable capsular polysaccharides and lipoproteins modify bacteriophage susceptibility, whereas in *Campylobacter jejuni*, *in vitro* phage-sensitivity assays and chicken infection experiments showed that phase-variable capsule modifications allow bacteria to avoid phage infection^31,32^.

More recently, single-cell transcriptomics examined the *B. fragilis* strain NCTC 9343 (the same strain used in this study) infected with the lytic phage Bf12P1 (which is genomically related to Barc2635 from this study). This work showed that individual-cell vulnerability to phage infection is influenced by multiple genetic loci, especially phase-variable CPS biosynthesis pathways and a predicted fimbriae operon^33^. This finding is directly relevant to the present work, which uses the *B*. *fragilis* phage Barc2635, a phage genomically related to the Bf12P1, because it provides direct evidence that phage exposure can separate infected and uninfected subpopulations in *B.* fragilis whose vulnerability is linked to phase-variable transcriptional states. Several studies now link DNA inversion-mediated phase variation to host inflammation and phage exposure *in vivo,* including Blandford et al., who showed that PSA promoter orientation differs in biopsies from patients with inflammatory bowel disease, supporting the relevance of phase-variable promoter states in host-associated disease contexts^13^. Carasso et al. subsequently showed that host inflammatory signals and bacteriophage presence reshape DNA inversion states across the gut microbiota, altering “ON”/”OFF” distributions at multiple loci and reprogramming community-level functions^11^. In *B. fragilis* specifically, Carasso et al. demonstrated that exposure to the lytic phage Barc2635 drives prolonged changes in DNA inversion states and supports long-term coexistence between phage and bacterial host rather than simple extinction^12^. Thus, *B. fragilis* provides a well-characterized *in vivo* system in which DNA inversion states, bacterial abundance, and phage abundance can be followed longitudinally during phage pressure. Together, these studies show that phase-variable surface and transcriptional states can affect phage susceptibility in specific bacterial systems, supporting the idea that lytic phage exposure may act as a selective force on particular bacterial cell states rather than uniformly across all cells.

At the same time, single-cell measurements and mathematical analyses^9^ have clarified how phase-variable systems generate population structure even without explicit phage selection. Lan et al. combined single-cell profiling of multiple invertible promoters in *Bacteroides* with stochastic computational modelling and demonstrated that differential inversion rates across promoters are a major determinant of bacterial population heterogeneity^9^. In their framework, each promoter is characterized by forward and reverse inversion rates, and the resulting Markovian switching process drives populations toward reproducible steady-state distributions of combinatorial phase-variable states. Despite the large number of possible “ON”/”OFF” combinations, the system converges over time to characteristic mixtures dictated by the inversion-rate matrix rather than by initial conditions. This work implies that the apparently high-dimensional phase-variable configuration landscape has an underlying dynamical structure that constrains which phenotypic configurations are actually occupied.

These experimental and modelling advances raise a central question: how does intrinsic phase-variable switching interact with extrinsic phage predation? In other words, when a multi-locus phase-variable system is exposed to a lytic phage, do multiple loci contribute independently, or is the response simplified into a smaller number of dominant susceptibility patterns? Previous theoretical and experimental work has primarily considered phase variation at only one or a few loci. For example, Sandhu et al. modelled *Campylobacter jejuni*–F336 dynamics in which simple-sequence-repeat-mediated “ON”/”OFF” switching at two capsular-modification loci, *cj1421* and *cj1422*, determines phage adsorption. The model showed that the balance between phage selection and fitness costs associated with resistant states can favour intermediate switching rates and phage persistence, but it remained restricted to a two-locus susceptibility space rather than the many-locus DNA-inversion architecture of *B. fragilis*^34^. More broadly, reciprocal evolution between bacterial resistance and phage infectivity can generate an arms race and fluctuating-selection dynamics that maintain phenotypic and genetic diversity^23^.

Phage–bacteria interactions are not static: bacterial populations can shift among susceptibility states, while phage abundance and infective capacity against different bacterial variants can also change over time. Clinical observations of changing bacterial susceptibility during treatment, together with experimental evidence that both bacterial susceptibility and phage infectivity can change over time, further illustrate the dynamic nature of phage–bacteria interactions^35,36^. Longitudinal analysis and mathematical modelling are therefore important for understanding and predicting these dynamics and for identifying candidate periods of increased bacterial susceptibility. However, existing models do not connect phage dynamics to bacterial longitudinal DNA inversion trajectories across multiple independently switching phase-variable regions. Thus, there remains a substantial gap for models that link phage dynamics to multi-locus phase variation in an *in vivo* bacterial–phage system such as *B. fragilis*–Barc2635, with broader relevance to bacterial persistence, phage–bacteria coexistence, and the prediction of time-dependent susceptibility and potential intervention windows. The scientific scope for the mathematical model is to determine how phage pressure organizes a high-dimensional set of reversible bacterial states over time, to infer changes in population-level susceptibility, to identify the loci and loci combinations that dominate these changes, and to generate testable predictions about bacterial persistence and susceptibility. This framework may also help identify candidate time windows in which bacterial populations are more susceptible to phage intervention, although its relevance to therapeutic timing will require validation.

In the present study, we address this gap by coupling a multi-locus phase-variable system to phage dynamics in the *B. fragilis* NCTC 9343-Barc2635 model. Our mathematical modelling is based on phage-bacteria *in-vivo* longitudinal interactions in gnotobiotic mouse datasets from Carasso et al., focusing on time-resolved trajectories of selected invertible regions during bacterial exposure to Barc2635^12^. We use an effective nonlinear ordinary differential equations (ODE) framework that represents “ON” and “OFF” bacterial orientation states for each invertible region, along with phage abundance, and an effective clearance term representing *in vivo* loss of bacteria and phage in the population-level model. Rather than modelling individual host processes explicitly, the host is represented through this effective clearance term, which captures the combined effects of host-associated clearance on both bacterial and phage populations. Fitted phage-associated parameters (e.g., adsorption rate, burst size and explicit phage decay parameter) are interpreted as model-derived susceptibility estimates and not as evidence that any specific promoter state is the physical Barc2635 receptor.

Our goals are to use this system as a quantitative framework and to ask how bacterial susceptibility to Barc2635 is organized across multiple phase-variable orientation states. By combining dynamical modelling, contribution-space analysis, and combinatorial scoring, we show that Barc2635-associated phase variation in *B. fragilis* is structured at multiple levels. Individual loci occupy a model-inferred susceptibility landscape, temporal dynamics are compressed into a small number of dominant contributors and organized into significant combinations. Thus, Barc2635-associated phase variation cannot be captured by a single susceptibility measurement; instead, it is organized through region-level differences, time-dependent contributions, and recurring multi-locus combinations.

## Results

### A longitudinal mouse analysis framework for modelling phage-associated phase variations in *B. fragilis*

To investigate how bacteriophage exposure affects phase-variable states in *Bacteroides fragilis*, we modelled the longitudinal germ-free mouse experiment from the *B. fragilis* NCTC 9343– Barc2635 study of Carasso et al^12^. This subset contained mice colonized with *B. fragilis* alone and mice colonized with *B. fragilis* infected with the Barc2635 bacteriophage. Fecal samples were collected at 13 time points from day 0 to day 49, producing 81 barcoded fecal samples for the modeling analysis (Supplementary table 1). Each sample was associated with phase-variable promoter orientation, bacterial abundance, and phage abundance measurements (Fig. 1A).

**Figure 1.**
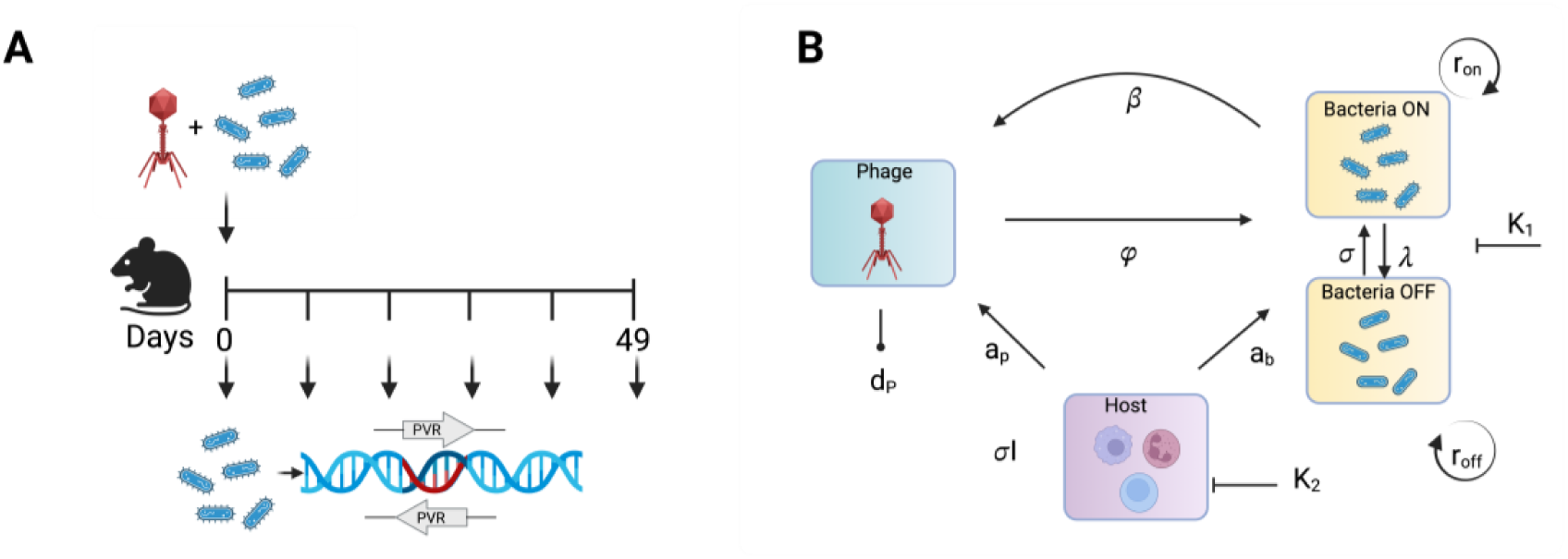
Experimental design and modeling framework. **(A)** Schematic of the longitudinal germ-free mouse experiment reanalyzed in this study. Germ-free mice were colonized with *Bacteroides fragilis* NCTC 9343 alone or with *B. fragilis* NCTC 9343 and bacteriophage Barc2635. The reanalysis used four longitudinal control mice and three longitudinal phage-exposed mice. Fecal samples were collected across 13 time points from day 0 to day 49 and used to quantify phase-variable “ON” fractions, bacterial abundance by colony forming units (CFU), and phage abundance by plaque forming units (PFU). **(B)** Conceptual model linking phase-variable “ON”/”OFF” bacterial orientation states, phage abundance, and an effective host-response or clearance term. Arrows indicate bacterial growth (r_on_ and r_off_), “ON”/”OFF” switching (sigma and gamma), effective phage interaction (phi and beta), phage decay (d_P_), and clearance of bacterial and phage compartments (a_b_ and a_p_), K1 and K2 are carrying capacities.

We focused on 18 invertible regions: seven capsular polysaccharide promoter loci, PSA, PSB, PSD, PSE, PSF, PSG, and PSH, together with 11 additional phase-variable regions, PVR1– PVR11. This design allowed us to ask whether Barc2635 exposure acts on individual invertible regions independently, or whether the multi-locus phase-variation system forms a coordinated susceptibility program.

We modeled each region using an “ON”/”OFF” dynamical framework coupled to phage abundance and a population-level clearance term that captures unmeasured *in vivo* loss of bacteria and phage (Fig. 1B). In this framework, promoter orientation is treated as a model- defined susceptibility-associated state, not as direct evidence that any specific locus is the physical receptor for Barc2635. Growth parameters were estimated from control trajectories, whereas phage-related parameters and effective switching rates were fit under phage exposure.

Detailed estimation, residual, and stochastic validation plots for each region are provided in Supplementary Fig. 1.

### Region-level modeling reveals a structured phage-susceptibility landscape

We first asked whether the 18 invertible regions differed in their effective association with Barc2635 dynamics. For each locus, we fit the dynamical model to longitudinal “ON”/”OFF” orientation, colony forming units (CFU), and plaque forming unit trajectories, then used the inferred parameters to define a region-level phage-sensitivity proxy. The resulting ranking showed a broad spectrum of effective susceptibility across loci (Fig. 2A). PVR5 had the highest phage-sensitivity score, followed by PSD, PVR11, PVR9, and PVR6. In contrast, PSG and PVR4 occupied the lowest end of the ranking. These results indicate that Barc2635-associated dynamics are not distributed uniformly across the phase-variable system.

**Figure 2.**
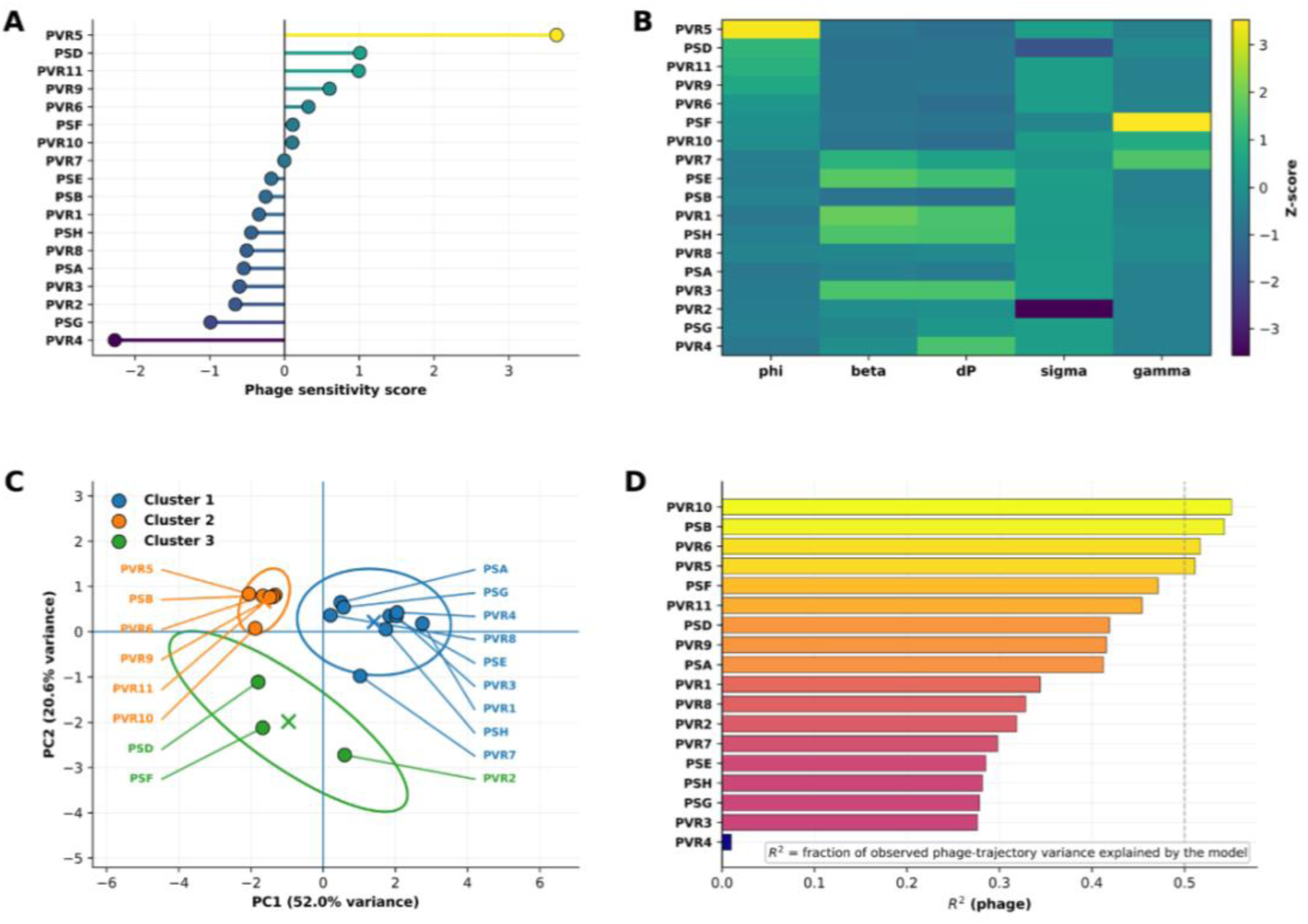
Region-level phage susceptibility landscape. **(A)** Ranked phage-sensitivity proxy scores across the 18 analyzed invertible regions. Higher values indicate stronger effective phage-associated susceptibility inferred from fitted model parameters. **(B)** Heatmap of standardized fitted phage-associated and switching parameters across regions. Columns show standardized values for effective phage interaction (phi), amplification (beta), phage decay (d_P_), and effective switching rates (sigma and gamma) under phage exposure. **(C)** Principal component analysis of standardized fitted parameters. Points represent phase variable regions, colors indicate k-means clusters with k = 3, ellipses show cluster dispersion, and crosses mark cluster centers. PC1 and PC2 explain 52.0% and 20.6% of the fitted-parameter variance, respectively. **(D)** Region-wise model fit to phage dynamics, quantified as the fraction of observed phage-trajectory variance explained by the model. The vertical dashed line marks a reference value of R² = 0.5. Variation in fit quality indicates that some loci are more strongly captured by the modeled phage-associated dynamics than others. Detailed per-region fits, residuals, and stochastic validation are shown in Supplementary Fig. 1.

The parameter heatmap showed that this ordering was not explained by a single fitted term (Fig. 2B). Instead, regions differed across multiple phage-associated and switching-related parameters, including effective phage interaction, amplification, decay, and “ON”/”OFF” transition behaviour. Thus, the region-level landscape reflects a coordinated parameter phenotype rather than one dominant variable.

To determine whether this parameter variation was high-dimensional or organized into simpler regimes, we performed PCA on the standardized fitted parameter matrix. The first two principal components captured most of the parameter-space structure, with PC1 explaining 52.0% and PC2 explaining 20.6% of the variance (Fig. 2C). K-means clustering separated the loci into three groups, indicating that the susceptibility landscape is structured into discrete dynamical classes rather than forming a completely diffuse continuum.

Model fit to phage dynamics varied across regions (Fig. 2D). Phage trajectories were best fitted for PVR10, PSB, PVR6 and PVR5, whereas the model captured little of the variation in the PVR4 phage trajectory.

### Contribution-space analysis identifies a temporally organized susceptibility program

The region-level model identified loci associated with Barc2635 sensitivity, but it did not directly show how these loci contributed to susceptibility over time. We therefore projected the fitted susceptibility weights onto the observed “ON”-fraction trajectories from the phage-exposed group. For each locus, this produced a time-resolved effective contribution that combines its model-inferred susceptibility potential with its realized promoter-orientation state. In this framework, a locus contributes strongly only when it has both a high susceptibility weight and sufficient “ON”-state representation during the time course. Consequently, contribution is not determined solely by the magnitude of the observed difference between phage and control groups, but by the combination of inferred susceptibility and the longitudinal orientation trajectory.

The resulting total modelled susceptibility trajectory tracked the major temporal organization of PFU and CFU dynamics (Fig. 3A). Susceptibility increased rapidly during the early phase, remained elevated and structured through the middle phase, and adapted during the late phase. Decomposition of the total signal revealed strong compression of the multi-locus system (Fig. 3B). Although 18 invertible regions were analyzed, a small subset accounted for most of the modeled susceptibility signal. PSF contributed persistently throughout much of the longitudinal time course, whereas PVR3, PVR5, PVR7, PSB, and several additional loci contributed predominantly during specific temporal windows. Thus, the high-dimensional phase-variable system is expressed over time through a smaller set of dominant effective contributors.

**Figure 3.**
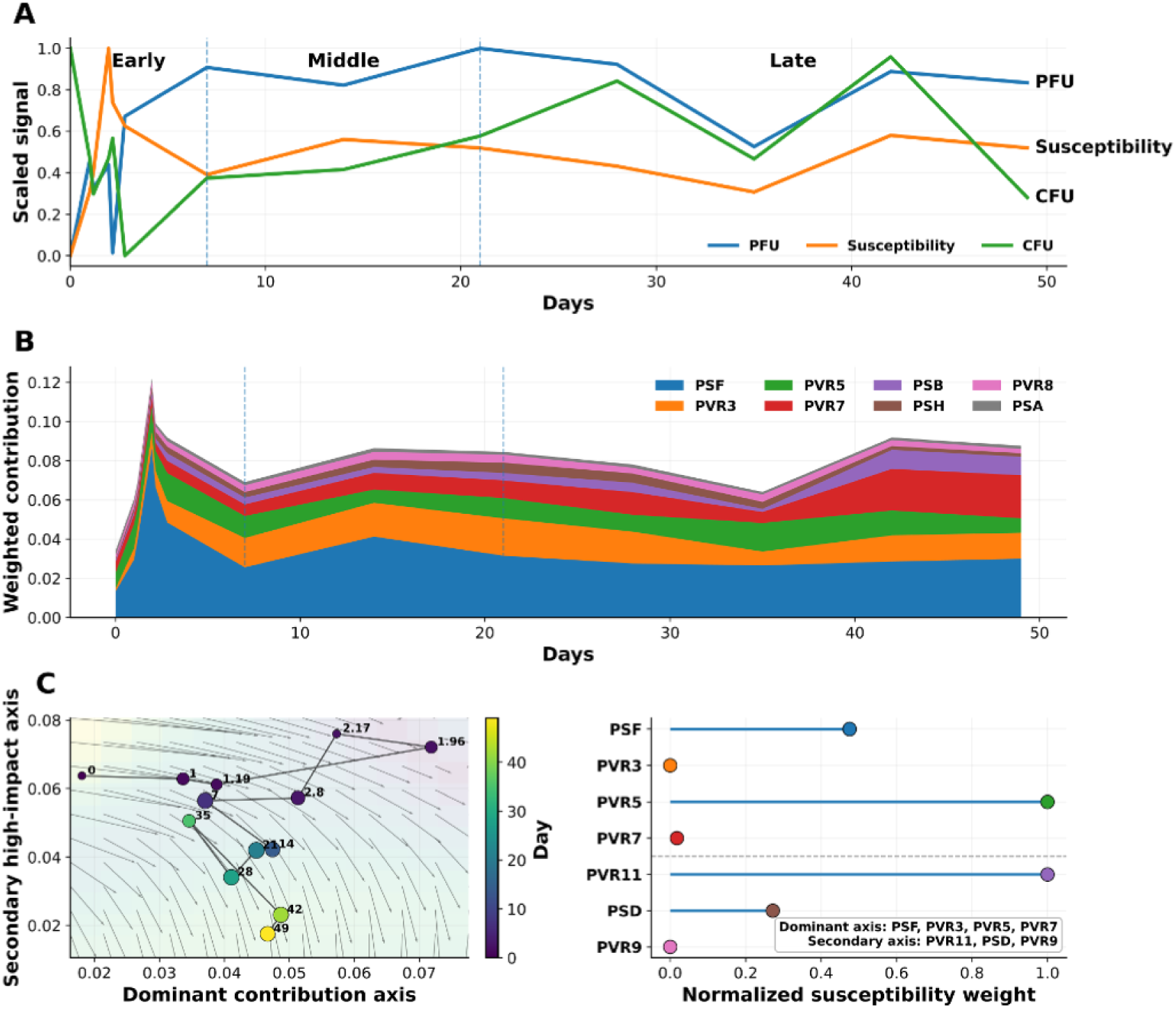
Dynamical organization of phage susceptibility in contribution space. **(A)** Temporal comparison of modeled susceptibility, phage abundance, and bacterial abundance. Total modeled susceptibility was computed from locus-specific susceptibility weights combined with observed phage-group ON fractions, then scaled together with PFU and CFU to a 0–1 range for visualization. Dashed vertical lines mark the early, middle, and late temporal phase boundaries used in the combinatorial analysis. **(B)** Stacked temporal decomposition of modeled susceptibility into dominant locus-level contributions. **(C)** Empirical flow field and temporal trajectory in two-dimensional contribution space. The left panel shows the trajectory through dominant and secondary contribution axes, with points colored by day and arrows indicating locally averaged direction of motion. The right panel shows the loci defining the dominant and secondary axes according to normalized susceptibility weight.

To visualize how this compressed susceptibility program evolved dynamically, we projected the temporal trajectory into a two-dimensional contribution space (Fig. 3C). This projection was not a PCA of fitted parameters. Instead, the axes were constructed from weighted combinations of selected loci. Here, susceptibility refers to the model-inferred ability of a locus to influence phage dynamics, quantified by its fitted susceptibility weight. However, a locus contributes to the observed trajectory only when this susceptibility potential is realized through sufficient representation of its “ON” orientation. Accordingly, we distinguish between susceptibility potential (high model-derived weight) and realized contribution (high weight combined with substantial “ON”-state occupancy over time). The dominant contribution axis was built from loci with the largest realized temporal contributions, whereas the secondary high-impact axis was built from loci with high susceptibility weights relative to their average “ON” fractions. The resulting trajectory followed an organized path rather than a diffuse scatter, and the empirical flow field showed coherent local directions of motion. The axis-defining loci clarified the distinction between persistent contribution and high-impact susceptibility potential. The dominant contribution axis was defined by PSF, PVR3, PVR5, and PVR7, whereas the secondary high-impact axis was defined by PVR11, PSD, and PVR9. PSF dominated the realized temporal signal because it contributed persistently across time. PVR5, by contrast, had the highest normalized susceptibility weight and strongly shaped the geometry of the dominant axis, but it contributed less consistently in the stacked temporal decomposition. Similarly, PVR11, PSD, and PVR9 contributed weakly on average but defined the secondary axis because their high weights allow transient activation to redirect the trajectory. Thus, Fig. 3B identifies the loci that dominate realized susceptibility through time, whereas Fig. 3C identifies the lower-dimensional axes that organize the trajectory.

### Combinatorial scoring reveals a PSF-centred susceptibility program across temporal phases

We next asked whether susceptibility arises from individual loci acting independently or from coordinated combinations of loci. To test this, we evaluated all pairwise, triple, and quadruple locus combinations separately within the early, middle, and late temporal phases. Each combination received a composite score based on two features: the summed susceptibility contribution of its member loci and the extent to which those loci were “ON” at the same time. Thus, a high score indicates that a combination contains loci with high model-inferred susceptibility weights and that these loci are temporally active together during that phase. Scores were normalized within each phase and combination size, so the values should be interpreted as relative rankings among candidate combinations, not as absolute probabilities of infection or direct measurements of receptor activity.

At the pairwise level, the strongest combinations were overwhelmingly PSF-centred (Fig. 4A). PSA + PSF and PSF + PVR4 were the highest-scoring early combinations. PSF + PVR3 peaked in the middle phase, while PSF + PVR4 remained strong across all phases. PSF + PVR7 became more prominent in the late phase. Thus, pairwise susceptibility is organized around a persistent PSF-containing core with phase-specific partners.

**Figure 4.**
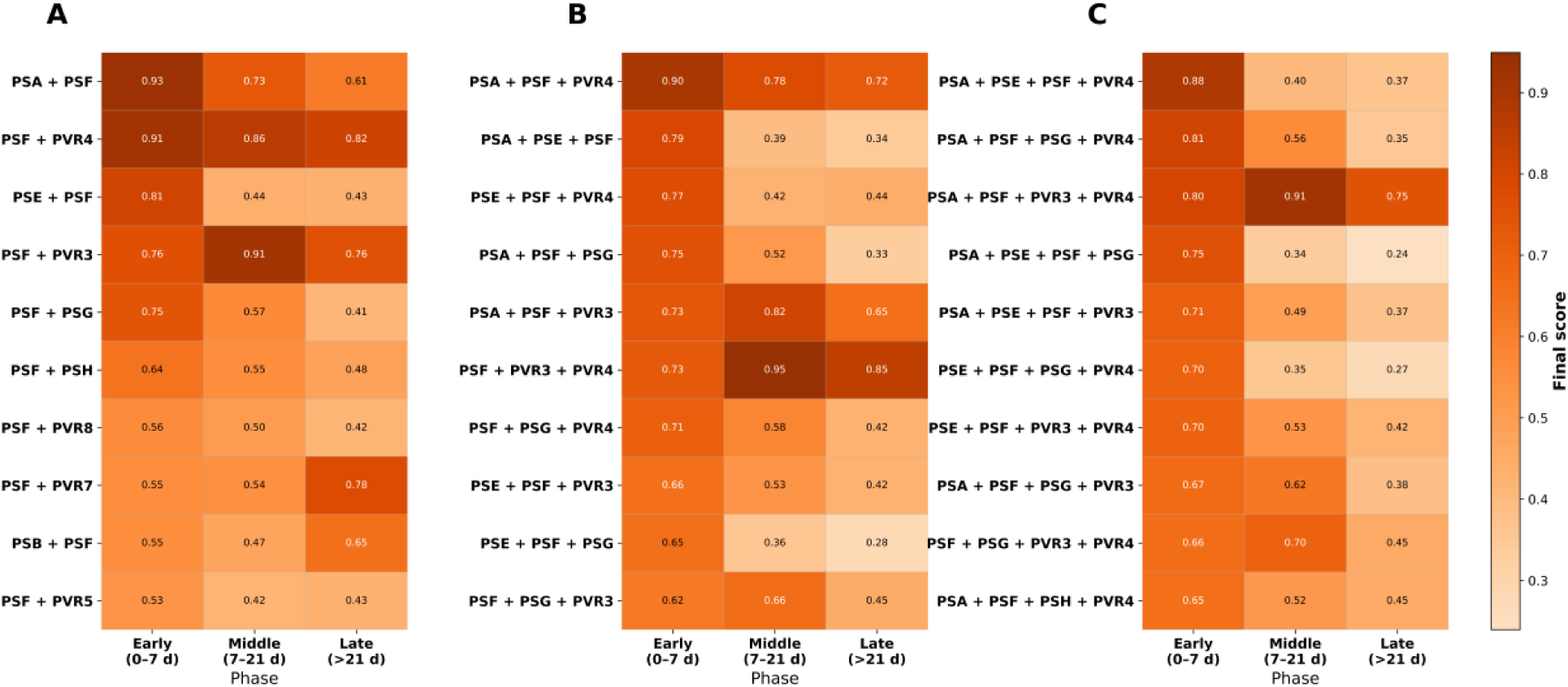
Combinatorial phage-susceptibility locus across continuous temporal phases. Heatmaps show the highest-scoring locus combinations for pairwise, triple, and quadruple sets across early, middle, and late temporal phases. Scores combine weighted additive contribution with a gated co-activation component, emphasizing combinations that are both susceptible and temporally coordinated. **(A)** Top pairwise combinations. **(B)** Top triple combinations. **(C)** Top quadruple combinations.

Triple combinations largely extended the same architecture (Fig. 4B). PSA + PSF + PVR4 was the strongest early triple, whereas PSF + PVR3 + PVR4 dominated the middle and late phases. Quadruple combinations introduced additional refinement but preserved a similar structure (Fig. 4C). PSA + PSE + PSF + PVR4 was strongest early, whereas PSA + PSF + PVR3 + PVR4 was the dominant middle-phase quadruple and remained strong in the late phase.

Mouse-level block bootstrap analysis supported the robustness of the inferred susceptibility organization (Figure 5). Across 5,000 mouse-level bootstrap replicates, PSF had median bootstrap rank 1 (Fig. 5A), with both bounds of the 95% bootstrap rank interval equal to rank 1 (Fig. 5B). PVR3, PVR5, PVR7, and PSB also remained in the top five in all replicates, whereas no other region entered the top five. Combination-level stability was phase-dependent. Early and middle phases showed strong PSF-centered robustness: PSA + PSF + PVR4 was the top early triple in 100% of replicates, PSF + PVR3 + PVR4 was the top middle triple in 100%, and PSA + PSF + PVR3 + PVR4 was the top middle quadruple in 100%. The late phase was more heterogeneous, with PSF + PVR4, PVR3 + PVR7, PSF + PVR3 + PVR4, and PSF/PVR3/PVR4/PVR7-containing quadruples alternating across bootstrap replicates.

**Figure 5.**
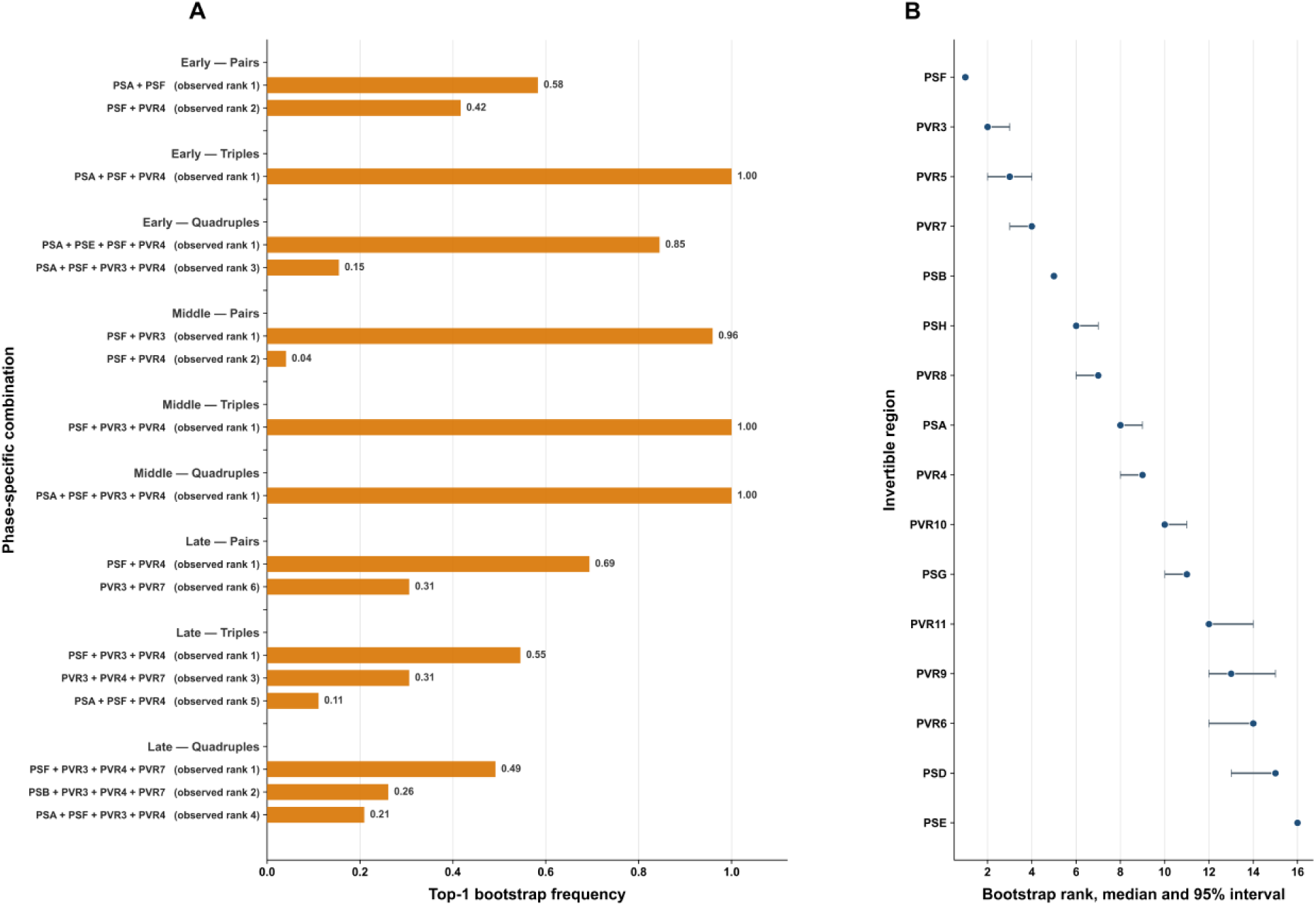
Mouse-level bootstrap robustness analysis. **(A)** Top-1 bootstrap frequency of phase-specific pairwise, triple, and quadruple locus combinations across 5,000 mouse-level block bootstrap replicates. Labels in parentheses indicate observed rank in the original dataset. **(B)** Median bootstrap rank and 95% bootstrap rank interval for retained regions based on mean effective contribution. Lower rank indicates stronger modeled contribution.

## Discussion

The biological foundation for this study was done by Carasso et al., who showed that exposure of *Bacteroides fragilis* NCTC 9343 to the lytic bacteriophage Barc2635 produces dynamic and prolonged changes in DNA inversion states during gut colonization^12^. In their gnotobiotic mouse system, phage exposure altered phase-variable regions, particularly promoter orientations associated with polysaccharide biosynthesis loci. These changes coincided with increased entropy in inversion ratios relative to control mice (with no phage exposure). Their study further showed that *B. fragilis* and Barc2635 can persist together over a multi-week time course. This bacteria-phage co-existence along with the dynamic bacterial DNA inversion alterations in response to phage exposure indicates that bacterial functionality can be reshaped without simple elimination of the bacterial host population^12^.

Here, we applied mathematical modelling on a defined longitudinal dataset from that study. Rather than asking whether exposure to Barc2635 leads to inversion state alterations, we asked how the measured inversion trajectories are organized when interpreted through a dynamical model of bacterial abundance, phage abundance, and phase-variable “ON”/”OFF” states. This distinction is important. While Carasso et al. demonstrated that phage-driven inversions are biologically meaningful and linked to long-term coexistence^12^, our analysis suggests that these inversion dynamics are not distributed randomly across loci but are organized into a smaller set of structured susceptibility patterns. Across three analysis layers, we found that Barc2635-associated phase variation is structured. First, region-level modeling revealed that the 18 analyzed invertible regions occupy a broad phage-susceptibility landscape. PVR5, PSD, PVR11, PVR9, and PVR6 ranked highly by the model-derived sensitivity proxy, whereas PVR4 and PSG ranked low. This ranking suggests that Barc2635 exposure is not equally coupled to all invertible regions. Instead, some loci have orientation trajectories and fitted parameter profiles that are more strongly aligned with longitudinal phage dynamics. High-ranking loci should be interpreted as candidate regions whose orientation trajectories and fitted parameter profiles are most strongly aligned with longitudinal phage dynamics, rather than as proven receptors or loci with an inferred "ON" state. Conversely, low-ranking loci, such as PSG, should not be interpreted as biologically irrelevant; rather, their behavior may be protective, context-specific, or poorly captured by a score that emphasizes sustained alignment with phage abundance over the full time course. The standardized parameter heatmap and PCA further showed that susceptibility is not controlled by a single fitted parameter; instead, loci differ in the fitted parameters describing effective phage susceptibility, phage amplification, phage decay, and switching behavior. This suggests that no single fitted parameter explains the inferred susceptibility landscape. Instead, high- and low-ranking loci reflect different combinations of fitted parameters, rather than simply differing in the proportion of "ON" states or the strength of phage interaction alone. Consequently, loci with similar overall susceptibility may achieve it through different combinations of dynamical mechanisms, and at the same time, loci with comparable promoter-orientation frequencies can differ substantially in their modeled susceptibility profiles. Therefore, high- and low-ranking loci should be interpreted as distinct dynamical strategies rather than simply as loci with more “ON”-oriented bacterial cells or stronger phage interaction alone. More broadly, these findings indicate that predicting bacterial susceptibility requires integrating multiple interacting processes, supporting the use of dynamical modeling instead of relying solely on static measurements of promoter orientation or individual fitted parameters.

Second, contribution-space analysis showed that the temporal response is compressed into a small subset of dominant contributors. Although 18 invertible regions were included, much of the modeled susceptibility signal was carried by PSF together with phase-dependent contributions from PVR3, PVR5, PVR7, PSB, and other loci. This result does not mean that PSF is highly “ON” in absolute terms, nor that it is the Barc2635 receptor. Instead, PSF is important because the contribution score combines fitted susceptibility weight with the realized “ON”- fraction trajectory. A promoter can remain low in absolute “ON” orientation yet contribute strongly if its “ON”-fraction trajectory remains consistently aligned with the temporal pattern of phage dynamics. Thus, loci differ not only in their inferred susceptibility weights but also in how their temporal “ON”-fraction trajectories align with the modelled phage dynamics. These conclusions were further supported by the bootstrap analysis, which confirmed a stable contribution hierarchy across resampled datasets. In particular, the recurrent dominance of PSF and the early-to-middle phase combinatorial patterns remained highly consistent, whereas greater variability emerged during the late phase, reflecting increased mouse-level heterogeneity at later time points. Thus, the bootstrap analysis confirms a stable contribution hierarchy and a robust early-to-middle PSF-centered combinatorial program, while indicating greater late-phase mouse-level variability. One possible biological explanation for the persistent importance of PSF is its proposed hierarchical role within the CPS regulatory network. Although our model does not explicitly represent regulatory interactions among promoters, the repeated dominance of PSF raises the hypothesis that changes at this locus may reflect broader reorganization of the capsular polysaccharide network. The PSF operon encodes the trans-locus inhibitor UpfZ, which can inhibit the expression of multiple other CPS operons, including PSA^12,21,37^. Thus, changes associated with PSF may reflect not only the state of this individual locus but also broader changes in capsule expression. However, because our model does not explicitly analyze cross-regulation among CPS operons, this remains open for future investigation.

Third, combinatorial scoring revealed that the modelled susceptibility program is not only low-dimensional but also organized around recurring multi-locus motifs. PSF-containing combinations dominated pairwise, triple, and quadruple rankings across early, middle, and late phases. PSA + PSF and PSF + PVR4 were prominent early, PSF + PVR3 and PSF + PVR3 + PVR4 became strongest in the middle phase, and PSF-containing combinations remained important until the later phases. This pattern suggests that increasing combinatorial complexity refines the same core program rather than replacing it with entirely different loci. PVR5 illustrates the distinction between individual susceptibility and coordinated susceptibility: it is highly weighted individually but largely absent from top combinatorial loci because the combinatorial score rewards temporal co-activation with other high-weight loci.

Recent single-cell transcriptomic work provides additional support for a PSF-relevant interpretation, while also highlighting an important difference between our model and direct short-term infection assays. Gupta et al. infected *B. fragilis* NCTC 9343 with Bf12P1, a lytic phage closely genetically related to Barc2635, and showed that single-cell vulnerability to phage infection was influenced by multiple loci, most prominently phase-variable CPS biosynthesis pathways and a predicted fimbrial operon^33^. Bf12P1 shares 89.0% nucleotide similarity with Barc2635 and readily infects and lyses *B. fragilis* NCTC 9343. In that study, PSF, PSB, and PSG capsule types were enriched among uninfected bacterial cells, and resistant cultures often selected PSF- or PSG-expressing states^33^. This convergence strengthens confidence in the biological relevance of PSF. Whereas Gupta et al. identified PSF-enriched resistant bacterial cells in a short-term infection assay^33^ and Carasso et al. observed prolonged PSF-associated changes *in vivo*^12^, our model places these observations into a quantitative longitudinal framework by showing that PSF consistently dominates the inferred susceptibility landscape despite its relatively low absolute “ON” abundance and in comparison with all other invertible regions (Supplementary Table 1).

In parallel, PSG highlights an important difference between the Gupta et al. single-cell infection results and our longitudinal modeling analysis: Gupta et al. identified PSG as a strongly protective capsule state against the Bf12P1 phage, whereas PSG ranked low in our Barc2635 model and did not appear among the dominant contribution-space or combinatorial results. Gupta et al. found that the “PSB/PSG” phase-locked strain resisted phage challenge, and PSG-’”ON” states were enriched in resistant isolates^33^. This difference may reflect the fact that our scoring framework emphasizes sustained temporal alignment with phage and bacterial abundance across a long *in vivo* time course, whereas PSG may act as a strong but transient or context-specific protective capsule state. Consistent with this interpretation, the longitudinal data of Carasso et al.^12^ showed an early increase in the PSG "ON" orientation after Barc2635 exposure, followed by a return toward predominantly "OFF" states, indicating a transient rather than a sustained response. Our framework is optimized to identify loci that shape longitudinal population dynamics rather than loci whose effects are restricted to brief or localized episodes.

Such transient effects may nevertheless be biologically important, particularly during the initial stages of phage exposure or under specific environmental conditions. Thus, PSG should not be dismissed as biologically irrelevant; rather, its absence from our top-ranked model outputs suggests that transient protection during phage exposure and persistent contribution to longitudinal population dynamics represent distinct biological phenomena.

This distinction is central to the interpretation of this study. A measured orientation shift identifies a promoter whose state changes under phage exposure. A fitted susceptibility weight identifies a locus whose trajectory helps explain phage dynamics under the model. A combinatorial score identifies loci whose activity is coordinated with other high-weight loci during a temporal phase. These quantities do not rank loci in the same order. The repeated prominence of PSF is particularly notable given its known position within the hierarchical CPS regulatory network of *B. fragilis*^21^, suggesting that the model may capture broader regulatory organization rather than simply changes in individual promoter orientation. The fact that PSF is central in contribution and combinatorial analyses despite low absolute “ON” orientation therefore supports a systems-level interpretation: phage-associated susceptibility depends on timing, persistence, and co-activation, not only on the magnitude of a single promoter’s “ON” fraction.

This also clarifies why our results should be viewed as a refinement of, rather than a contradiction to the entropy/bet-hedging interpretation from Carasso et al.^12^: In that study, increased entropy reflected a broader distribution of bacterial cells across phase-variable “ON”/”OFF” states under phage exposure, which was interpreted as a bet-hedging strategy. The rationale is that a more diverse population is less likely to be uniformly susceptible to a single selective pressure, increasing the probability that at least some bacterial subpopulations will survive and repopulate when conditions change. Our analysis asks which parts of that diverse inversion landscape are most aligned with a modelled response to phage exposure. These two views can coexist because phage exposure may increase population-level phase-variable diversity, while the phage-associated component of that diversity is concentrated into a smaller number of weighted temporal contributors and recurring combinations.

The additional PVRs also help explain why a multi-locus framework is necessary. PVR3 repeatedly appears as a strong partner of PSF, especially in middle-phase combinations, and its genomic architecture is more complex than a simple promoter switch because it contains a long invertible cassette with predicted internal coding potential. PVR4 frequently appears in top combinations, suggesting that its importance lies in the combinatorial context rather than in its individual contribution. PVR7 becomes more prominent in later phases, consistent with continuing reorganization during long-term coexistence. PVR11, PSD, and PVR9 define the secondary high-impact axis in the contribution space. This suggests that some loci can strongly influence the modeled trajectory even if their effects are confined to particular time windows. In contrast, loci that look striking in the measured inversion data, such as PSG or PVR1 in the source study, may not rank highly in our mathematical model contribution analysis. This can happen if their changes are short-lived, do not align well with phage amplification, or do not occur together with other high-weight loci.

More generally, the architectural notes show that most PVRs share promoter-switch organization but vary in cassette length, inverted-repeat properties, and coding potential. This leads to the conclusion that no single architectural feature fully explains the inferred susceptibility landscape. An important conceptual advancement of this work is that the mathematical model does not replace the experimental observations but organizes them into a predictive dynamical framework. Previous studies, including the work of Carasso et al. and Gupta et al., demonstrated that phage exposure is associated with changes in phase-variable states and that particular promoter configurations can influence susceptibility to infection. Our results do not argue that phase variation itself becomes deterministic or non-stochastic under phage pressure. Rather, they suggest that stochastic switching is filtered through phage selection into reproducible population-level patterns that can be quantified mathematically. The model further distinguishes loci with transient changes from those that exert persistent influence on longitudinal phage dynamics. It also identifies coordinated multi-locus programs that are not apparent from individual promoter trajectories alone, and generates quantitative, experimentally testable predictions about the temporal organization of susceptibility. More broadly, this framework provides a general strategy for integrating longitudinal phase-variation data with dynamical modelling in other host-bacteria–phage systems. Future applications of the mathematical model could be used to improve the understanding of when bacterial populations transition into phase-variable states associated with increased or decreased susceptibility to phage infection.

Overall, Carasso et al. showed that Barc2635 drives DNA inversions that shape *B. fragilis* functionality and enable long-term phage–bacteria coexistence. Our analysis extends that finding by showing that the response has quantitative structure. Individual loci occupy a model- inferred susceptibility landscape, and the temporal response is compressed into a few dominant contributors. The highest-scoring combinations repeatedly converge on a PSF-centered, phase-dependent program, resulting in the conclusion that one promoter alone does not determine phage susceptibility, but that phage exposure reveals a constrained multi-locus organization in which low-abundance, persistent, and temporally aligned phase-variable states can have disproportionate dynamical influence. Importantly, this organization is not directly evident from the raw inversion trajectories alone but emerges only after integrating the dynamics of longitudinal orientation and phage population through the mathematical framework. This highlights the added value of quantitative modeling for distinguishing persistent drivers from transient responses and for revealing coordinated susceptibility programs that are otherwise difficult to identify experimentally.

## Limitations

Several limitations are important to acknowledge. First, the analysis uses bulk fecal measurements, so it does not directly observe single-cell capsule expression, phage adsorption, or infection outcomes for each orientation state. Second, the inferred susceptibility weights are model-derived and should not be interpreted as receptor identity. Third, model validation is primarily internal: per-locus fitting, residual analysis, stochastic simulations, and bootstrap analysis support consistency of the framework, but they do not replace experimental perturbation. Fourth, some high-scoring loci, including low-amplitude or near-static PVRs, should be treated as hypotheses requiring additional identifiability and perturbation analysis, because fitted weights can be less constrained when the underlying orientation trajectory changes only weakly. Fifth, the precise ordering of combinatorial scores depends on the chosen scoring framework, including the predefined weighting of peak versus mean and additive versus gated components. However, the repeated prominence of PSF across contribution-space analysis, multiple combination sizes, temporal phases, and mouse-level bootstrap resampling suggests that the broader PSF-centered organization is not solely attributable to a single combinatorial ranking. Sixth, the effective host-response or clearance term in the model should also be interpreted conservatively. It represents population-level clearance and other loss processes rather than a directly measured immune pathway. Its role is to constrain the coupled bacteria– phage dynamics and prevent biologically unrealistic trajectories, rather than to assign a specific immunological mechanism to PSF, PVR3, PVR5, or any other locus. In germ-free mice, which are not immune-free, such processes may include host-associated clearance, gut transit, non-specific phage loss, mucus-associated effects, and other unmeasured *in vivo* factors that shape bacterial and phage trajectories. In germ-free mice, which are not immune-free, such processes may include host-associated clearance, gut transit, non-specific phage loss, mucus-associated effects, and other unmeasured *in vivo* factors that shape bacterial and phage trajectories. Seventh, the temporal resolution of the available longitudinal dataset limits the ability to resolve the precise ordering of regulatory events. Although the current sampling, together with the modelling framework, is sufficient to distinguish sustained from transient contributors and to identify recurring temporal patterns, it does not determine whether changes in one phase-variable region precede and contribute to changes in downstream loci through the known CPS regulatory hierarchy. Such regulatory cascades remain mechanistic hypotheses beyond the scope of the present population-level analysis.

These limitations point to direct experimental tests. Locked-orientation mutants for PSF, PVR3, PVR4, PVR5, PVR7, PSD, PVR9, and PVR11 would test whether the inferred contributors causally alter Barc2635 susceptibility. Phage adsorption assays against defined capsule or PVR states would distinguish direct receptor effects from indirect population-level associations. Single-cell measurements that jointly capture promoter orientation, capsule expression, and phage binding would test whether low-frequency PSF-”ON” bacterial cells are disproportionately involved in phage interaction or coexistence dynamics. The Gupta et al. ^33^ study shows how powerful this single-cell approach can be, but an analogous experiment with Barc2635 in the *in vivo*-relevant context would be needed to test whether the PSF- and PSG- associated patterns observed with the Bf12P1 phage generalize to the Barc2635 phage. Finally, repeating the same modeling framework across additional phages could determine whether the PSF-centered organization is specific to Barc2635 or reflects a broader principle of *B. fragilis* phase-variable susceptibility.

## Methods

### Source data and reanalysis subset

This study reanalyzed a subset of the longitudinal germ-free mouse experiment reported by Carasso et al.^12^. In the original experiment, germ-free mice were colonized with *Bacteroides fragilis* NCTC 9343 either alone or together with the lytic bacteriophage Barc2635, and fecal samples were collected longitudinally for bacterial abundance, phage abundance, and DNA inversion-state analysis. The present study did not repeat the animal experiment, bacterial culture, phage propagation, sample collection, DNA extraction, or sequencing procedures. These experimental details are described in the source study^12^ and in Supplementary table 1.

The reanalysis table contained 81 barcoded fecal samples from control and phage-exposed groups. The control group contained four longitudinal mice colonized with *B. fragilis* alone, and the phage-exposed group contained three longitudinal mice colonized with *B. fragilis* and Barc2635. Samples were collected at 13 time points: days 0, 1, 1.19, 1.96, 2.17, 2.8, 7, 14, 21, 28, 35, 42, and 49.

For each sample, the table contained phase-variable “ON”-fraction measurements, bacterial abundance measured as colony-forming units (CFU), and phage abundance measured as plaque-forming units (PFU). Each barcode corresponded to one mouse fecal sample. For model fitting and figure generation, per-sample values were summarized as condition-by-time trajectories for each region.

### Phase-variable regions analyzed

Eighteen invertible regions were included in the analysis. These comprised seven capsular polysaccharide promoter loci, PSA, PSB, PSD, PSE, PSF, PSG, and PSH, together with 11 additional phase-variable regions, PVR1-PVR11. These loci were selected because they were monitored in the source dataset and included both canonical polysaccharide-associated promoter regions and additional invertible regions with phage-associated orientation changes.

For each locus *i*, the measured “ON” fraction at time *t* was denoted *x_i_*(*t*). Bacterial abundance was denoted *N*(*t*), and phage abundance was denoted *P*(*t*). For fitting the full “ON”/”OFF” model, observed “ON” and “OFF” bacterial abundances were reconstructed as

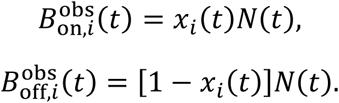

Phase-variable orientation measurements were taken from the processed experimental data generated from the source study. Large-scale detection and quantification of invertible promoter states in bacterial genomes is commonly performed using approaches such as PhaseFinder^7^.

### Full dynamical model of phage-phase-variation dynamics

For each invertible region, we modeled the bacterial population as two effective subpopulations corresponding to the “ON” and “OFF” orientation states of that region. The model tracked four variables: “ON” bacteria, “OFF” bacteria, free phage, and an effective host/clearance variable:

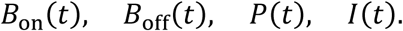

The full model was

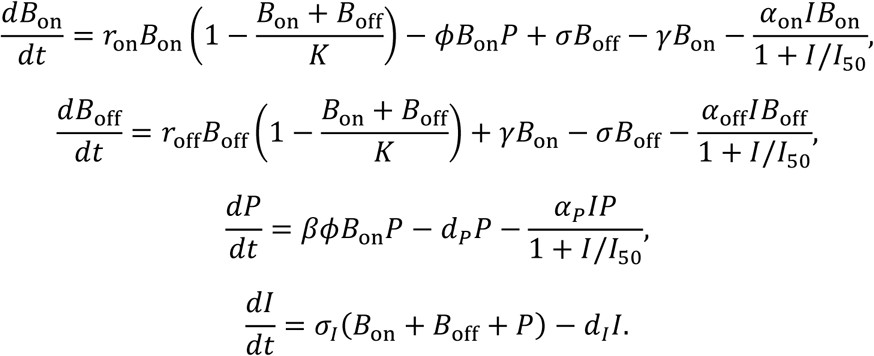

Here, *r*_on_ and *r*_off_ are growth rates of the “ON” and “OFF” bacterial subpopulations, *K* is the carrying capacity, *σ* is the effective “OFF”-to-”ON” switching rate, *γ* is the effective “ON”-to- “OFF” switching rate, *φ* is the effective phage-interaction parameter, *β* is the phage amplification factor, and *d_P_* is the phage decay rate. The parameters *α*_on_, *α*_off_, and *α_P_* describe effective clearance of “ON” bacteria, “OFF” bacteria, and phage, respectively. The parameter *I*_50_ controls saturation of the clearance term, *σ_I_* controls stimulation of the effective clearance variable, and *d_I_* controls its decay.

The phage-associated interaction term was parameterized through the “ON” compartment as a working modeling assumption for each locus. This does not establish that “ON”-oriented bacterial cells are the only infectable state, nor does it identify any specific invertible region as the physical receptor for Barc2635. Throughout the manuscript, fitted phage-interaction parameters are interpreted as effective phage-associated susceptibility terms.

### Effective host/clearance term

The variable *I*(*t*) was included to represent unmeasured host and environmental processes that can constrain bacterial and phage abundance *in vivo*. Although the mice were germ-free, they were not free of host physiology, gut transit, innate clearance, mucus-associated effects, or other loss processes. Therefore, *I*(*t*) was treated as an effective host/clearance variable rather than a measured immune pathway.

Clearance was modeled with a saturating functional form,

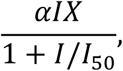

where *X* represents the relevant bacterial or phage compartment. This formulation approximates mass-action-like clearance at low *I* while preventing unrealistically strong elimination at high *I*. The term is phenomenological and should not be interpreted as evidence for a specific immune mechanism.

### Parameter estimation strategy

Parameter estimation was performed separately for each invertible region. The fitting strategy was designed to separate growth-related constraints estimated from control mice from phage-associated dynamics estimated under Barc2635 exposure.

### Control-condition fitting

Control trajectories, in which mice were colonized with *B. fragilis* without Barc2635, were first used to estimate bacterial growth behavior. For each locus, the “ON”-fraction trajectory and

CFU trajectory were used to reconstruct 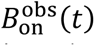(*t*) and 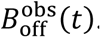(*t*). Growth parameters *r*_on_ and *r_off_* were estimated from the control trajectories. The carrying capacity *K* was fixed to a shared control-derived value across regions.

Control-condition switching parameters were also estimated for descriptive comparison, but they were not fixed during phage-condition fitting. This distinction is important because switching rates in the phage-exposed model were treated as effective condition-specific parameters.

### Phage-condition fitting

Phage-exposed trajectories were fitted with *r*_on_, *r*_off_, and *K* constrained by the control-condition fit. In contrast, phage-related parameters *φ*, *β*, and *d_P_*, together with effective switching rates *σ* and *γ*, were estimated jointly from the phage-exposed trajectories. This allowed switching dynamics to differ under phage exposure while preventing growth parameters from being re-fit to absorb phage-associated effects.

This strategy supports the interpretation that *σ* and *γ* in the phage model represent effective switching dynamics under phage exposure, not fixed intrinsic properties of promoter architecture.

### Objective function and numerical optimization

Model fitting was performed by nonlinear least squares with multi-start initialization. For each region, the optimization was repeated from multiple randomly initialized parameter sets, and the best-fitting solution was selected according to the minimized residual sum of squares.

Residuals were computed on log-transformed abundance values to reduce dominance by large absolute differences at high population sizes. Bacterial and phage abundances were transformed as

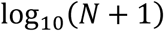

and

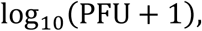

respectively. For ON/OFF bacterial compartments, observed values were reconstructed from the measured “ON” fraction and CFU abundance before log transformation. Parameter bounds and initialization ranges used in the optimization are provided in the analysis scripts and parameter tables.

### Model validation and fit assessment

Model performance was evaluated using deterministic fits to the observed longitudinal trajectories. For each region, observed and predicted trajectories were compared for “ON” bacteria, “OFF” bacteria, and phage abundance. Phage fit quality was summarized using the coefficient of determination,

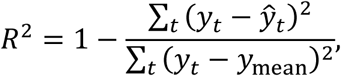

where *y_t_* is the observed log_10_(PFU + 1), *y^_t_* is the model-predicted value, and *y*_mean_ is the mean observed value. Region-specific fit quality was used to evaluate how strongly each locus was captured by the modeled phage-associated dynamics.

Detailed deterministic fits, residuals, fitted trajectories, and stochastic validation plots for each of the 18 regions are provided in Supplementary Fig. 1.

### Stochastic validation

Stochastic validation was used to assess whether the deterministic model trajectories were robust to effective variability around the fitted solution. Noise amplitudes were estimated from the dispersion of residuals between observed and predicted phage trajectories on the log_10_(PFU + 1) scale. These noise terms therefore combine measurement variability and unexplained dynamical variability and should not be interpreted as direct measurements of biological noise.

Stochastic trajectories were generated by applying multiplicative perturbations around the deterministic phage trajectories. Where implemented dynamically, perturbations were applied as diagonal noise terms around the deterministic vector field. These simulations were used as internal validation of model consistency and robustness, not as out-of-sample predictive validation.

### Region-level susceptibility landscape

For each region, fitted phage-exposed parameters were assembled into a feature matrix,

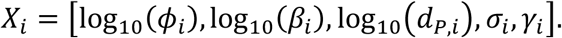

Each feature was standardized across regions before downstream analysis. Standardized parameters were visualized as a heatmap to compare the balance of phage interaction, phage amplification, phage decay, and effective switching behavior across loci.

A composite phage-sensitivity proxy score was used to rank regions in the region-level susceptibility landscape. This score was derived from fitted phage-associated and effective switching parameters and was used only as a relative ranking measure. The exact implementation used for the plotted score is provided in the analysis scripts. Downstream contribution-space and combinatorial analyses used fitted amplification-to-decay weights from the forced-phage model rather than this proxy score.

### Principal component analysis and clustering

Principal component analysis was applied to the standardized fitted-parameter matrix to visualize the dominant axes of variation among the 18 invertible regions. PCA was used as an exploratory dimensionality-reduction method and was not used as the sole basis for biological interpretation.

K-means clustering with *k* = 3 was applied in principal-component space to identify broad dynamical regimes among the regions. Cluster centroids and covariance ellipses were plotted for visualization. The number of clusters was chosen to summarize the observed high-, intermediate-, and low-susceptibility parameter regimes.

### Reduced forced-phage model

To isolate phage amplification from the full coupled host-phage model, we constructed a reduced forced-phage model in which the experimentally observed “ON” fraction and bacterial abundance were treated as time-dependent inputs. For each locus *i*, the observed phage-group “ON” fraction *x_i_*(*t*) was interpolated across time points, and bacterial abundance was scaled to obtain *Ñ*(*t*). Scaled phage abundance *P̃*(*t*) was used as a relative infection-pressure term in the susceptibility equation.

The reduced model was

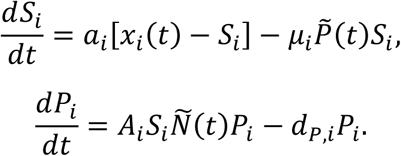

Here, *S_i_*(*t*) is an effective susceptibility state for locus *i*, *P_i_*(*t*) is modeled phage abundance, *a_i_* describes relaxation of susceptibility toward the observed “ON”-fraction trajectory, *μ_i_* describes phage-associated depletion of susceptibility, *A_i_* is the effective phage-amplification strength, and *d_P_*_,*i*_ is the phage decay parameter.

The forced-phage model was fitted by nonlinear least squares with multi-start initialization, minimizing residuals between observed and predicted log_10_(PFU + 1). This model was used to derive susceptibility weights for contribution-space and combinatorial analyses.

### Susceptibility weights

For each retained locus, a normalized susceptibility weight was computed from the forced-phage model as

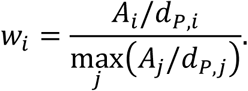

Only loci with positive amplification-to-decay ratios and positive phage-fit quality were retained for contribution-space analysis. The weight *w_i_* measures effective phage amplification relative to decay and should be interpreted as a model-derived susceptibility weight, not as a direct receptor measurement.

### Contribution-space analysis

To connect fitted phage sensitivity with observed temporal phase variation, the effective contribution of each locus was defined as

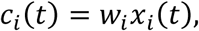

where *x_i_*(*t*) is the phage-group “ON” fraction and *w_i_* is the normalized susceptibility weight. Total modeled susceptibility was computed as

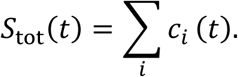

For visualization, total susceptibility, PFU, and CFU trajectories were min-max scaled to the interval 0-1:

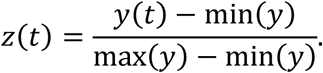

This scaling was used only for overlay visualization and did not affect contribution calculations.

Dominant temporal contributors were identified by ranking loci according to their mean effective contribution,

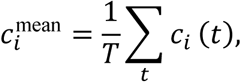

and peak effective contribution,

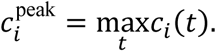

A two-dimensional contribution space was then constructed. The dominant contribution axis was defined from loci with the largest realized temporal contributions. The secondary high-impact axis was defined from loci with high susceptibility weights relative to their average “ON” fractions.

For the final contribution-space figure, the dominant axis was defined by PSF, PVR3, PVR5, and PVR7, and the secondary high-impact axis was defined by PVR11, PSD, and PVR9. If *D* denotes the dominant-axis loci and *R* denotes the secondary-axis loci, the axes were computed as

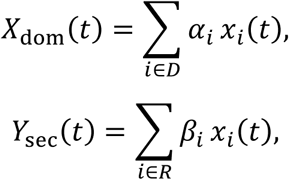

where *α_i_* and *β_i_* are normalized weights within each selected set.

### Empirical flow field in contribution space

Temporal motion in contribution space was estimated from consecutive time-point differences:

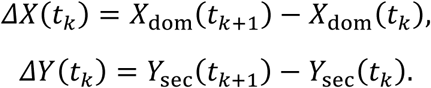

These local displacement vectors were smoothed across a grid in contribution space using distance-weighted averaging. The resulting empirical vector field was plotted together with the observed temporal trajectory, with points colored by sampling day. This analysis was used to visualize whether susceptibility dynamics followed constrained temporal paths rather than a diffuse distribution across contribution space.

### Temporal phase definitions

For contribution and combinatorial analyses, the time course was divided into three continuous temporal phases:

- Early: 0-7 days
- Intermediate: 7-21 days
- Late: >21 days

These boundaries correspond to the early post-colonization and phage-response interval, an intermediate coexistence interval, and a late persistence interval. The same phase definitions were used for pairwise, triple, and quadruple combinatorial scoring.

### Combinatorial scoring of locus sets

To quantify higher-order locus combinations, all combinations of size *k* = 2, *k* = 3, and *k* = 4 were evaluated. For each locus *i*, the normalized susceptibility weight *W̃_i_* was defined from the forced-phage amplification-to-decay ratio:

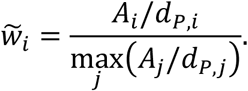

For a combination *C*, two complementary time-dependent scores were computed. The additive score was

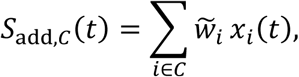

which captures cumulative weighted activity across loci. The gated score was

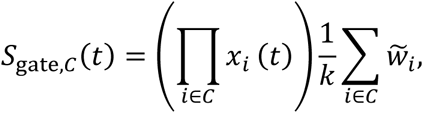

which emphasizes coordinated co-activation by penalizing combinations in which one or more loci are inactive.

Within each temporal phase, additive and gated scores were independently normalized across combinations:

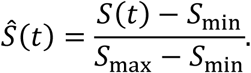

For each combination and phase, peak and mean values were then computed for both normalized scores. The final composite score was

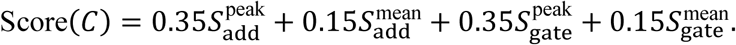

These weights were chosen a priori to balance two features of the combinatorial response. First, additive and gated components were given equal total weight, with each contributing 50% of the final score. The additive component captures the cumulative weighted activity of the loci in a combination, whereas the gated component rewards simultaneous co-activation and penalizes combinations in which one or more loci are inactive. Second, peak values were given greater weight than phase means, with 70% of the final score assigned to peak behavior and 30% to mean behavior. This choice reflects the expectation that phage-associated selection may act through short, phase-specific susceptibility windows, while still retaining information about combinations that remain active throughout a phase. The composite score was used only for relative ranking of combinations within each temporal phase and was not fitted to the data or optimized to favor any specific locus. Combinations were ranked within each phase according to this score, and the highest scoring pairwise, triple, and quadruple combinations were visualized as heatmaps using a shared color scale.

### Mouse-level bootstrap and rank-stability analysis

To assess whether the inferred contribution and combinatorial rankings were robust to individual phage-exposed animals, we performed mouse-level block bootstrap analysis. Each bootstrap replicates sampled phage-exposed mice with replacement while preserving each selected mouse’s full longitudinal trajectory. For each replicate, phage-group “ON”-fraction trajectories were recomputed for each retained invertible region and combined with the fixed model-derived susceptibility weights from the reduced forced-phage model. The susceptibility weights were derived from the fitted amplification-to-decay ratio and were interpreted as model-derived susceptibility weights rather than direct receptor measurements. Loci with zero or excluded susceptibility weight were not included in the weighted contribution and combinatorial ranking analysis.

Bootstrap analysis^38^ was performed using 5,000 replicates with random seed 12345. The retained regions were PSA, PSB, PSD, PSE, PSF, PSG, PSH, and PVR3–PVR11. PVR1 and PVR2 were excluded from weighted ranking because they did not pass the retained-weight criteria in the forced-phage analysis. PSA and PSE were orientation-flipped for the combinatorial scoring, consistent with the original combination analysis. For each replicate, we recomputed region-level effective contribution scores, region ranks, and all pairwise, triple, and quadruple combination scores across early, middle, and late temporal phases. Stability was summarized as median bootstrap rank, 95% bootstrap rank interval, top-one frequency, top-three frequency, and top-five frequency. Because only three phage-exposed mice were available for resampling, these bootstrap frequencies were interpreted as animal-level robustness measures rather than formal population-level p-values.

### Software and reproducibility

Model fitting, numerical integration, PCA, clustering, contribution-space construction, combinatorial scoring, and figure generation were performed using custom MATLAB and Python scripts. MATLAB R2018a and Python 3.10.14 were used for the analyses. Statistical analyses and figure generation were performed in Python 3.10.14 using NumPy^39^ 1.26.4, pandas^40^ 2.3.0, SciPy^41^ 1.15.2, and Matplotlib^42^ 3.10.1. Nonlinear least-squares optimization was performed with multi-start initialization, and numerical ODE integration was performed with stiff solvers where required.

All scripts required to reproduce preprocessing, model fitting, validation, contribution-space analysis and combinatorial scoring are available at: https://github.com/Julia-Be-sudo/Phase_Modeling

## Data availability

The experimental data reanalyzed in this study were generated in the previously published *Bacteroides fragilis* NCTC 9343–Barc2635 germ-free mouse study by Carasso et al.^12^. No new animal experiments, sequencing experiments, or phage-infection experiments were performed for the present study. The processed modeling input table used here, including sample identifiers, experimental group, sampling time, ON-fraction measurements, CFU values, and PFU values, is provided as Supplementary Table 1. Fitted model parameters, model-validation metrics, susceptibility weights, contribution scores, combinatorial scores, bootstrap outputs, and PVR genomic annotations are provided as Supplementary Table 2 and the Source Data files accompanying this manuscript. Source data underlying all main and supplementary figures are provided with this paper. Raw experimental and sequencing data from the original mouse experiment are available as described in Carasso et al.^12^.

## Acknowledgements

We thank Shaqed Carasso, Roni Keshet-David and Jia Zhang for generating and publishing the *B. fragilis*–Barc2635 longitudinal experimental dataset reanalyzed in this study, and for the foundational experimental work that made the mathematical modeling and this study possible. We thank Rachel Herren for critical reading of the manuscript and constructive comments. Finally, we thank the Geva-Zatorsky lab members for fruitful discussions and contributions.

## Funding

This project received funding from the Technion Institute of Technology, “Keren haNasi,” Cathedra, the Rappaport Technion Integrated Cancer Center, the Alon Fellowship for Outstanding Young Researchers, the Uzi & Michal Halevy Fund For Innovative Applied Engineering Research, the Israeli Science Foundation (1175/25), CIFAR (grant FL-000969/FL-001245/FL-001381), and the European Union (ERC, ExtractABact, 101078712). Views and opinions expressed are, however, those of the author(s) only and do not necessarily reflect those of the European Union or the European Research Council Executive Agency. Neither the European Union nor the granting authority can be held responsible for them. N.G.-Z. is a CIFAR fellow in the Humans & the Microbiome Program, a Kavli fellow, and a Horev Fellow (Taub Foundation).

## Notes

### Competing Interest Statement

The authors have declared no competing interest.

