## Supplemental Figure 1 for "Mathematical Modelling of Bacterial DNA Inversion Dynamics Uncovers an Organized Multi-Locus Response to Phage Predation in *Bacteroides fragilis*"

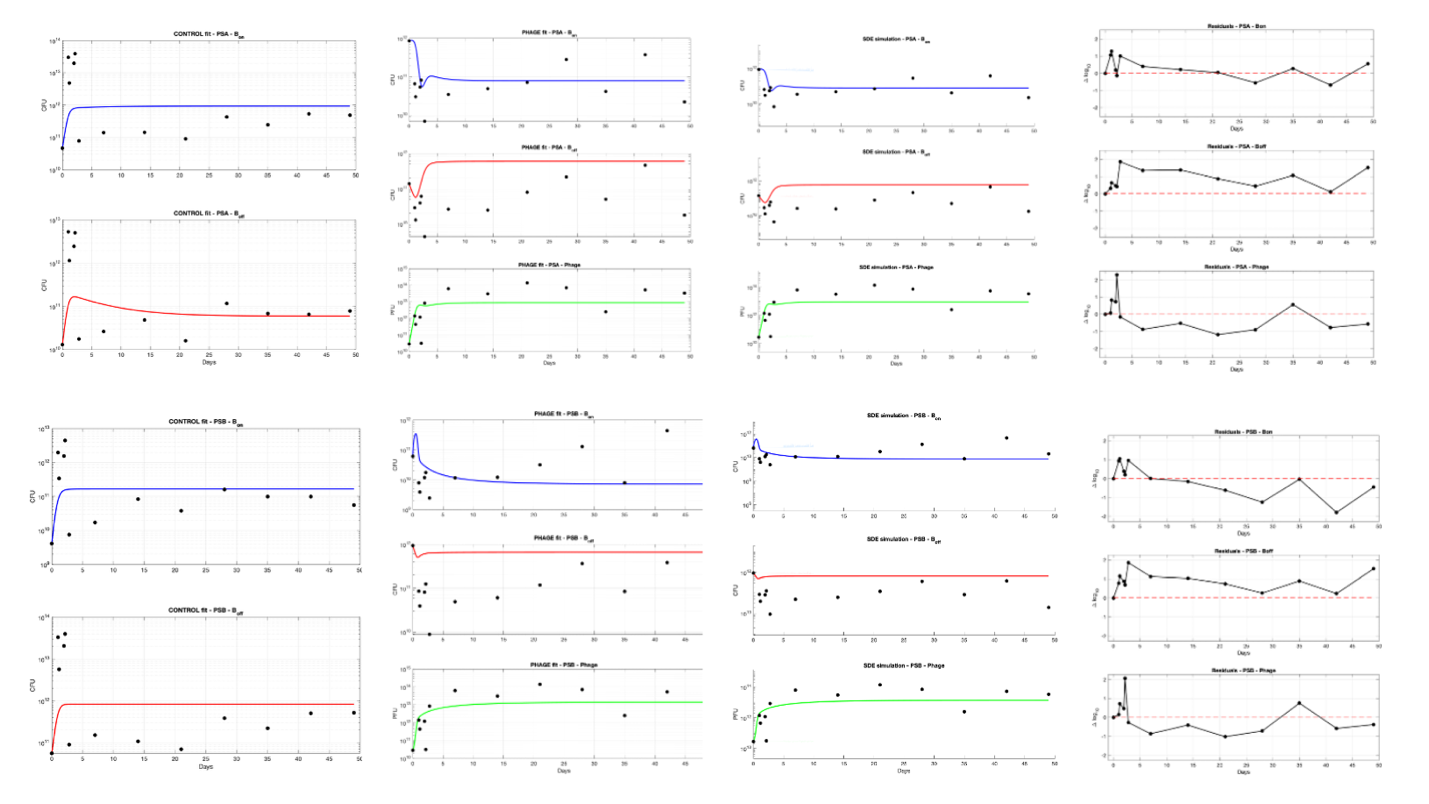

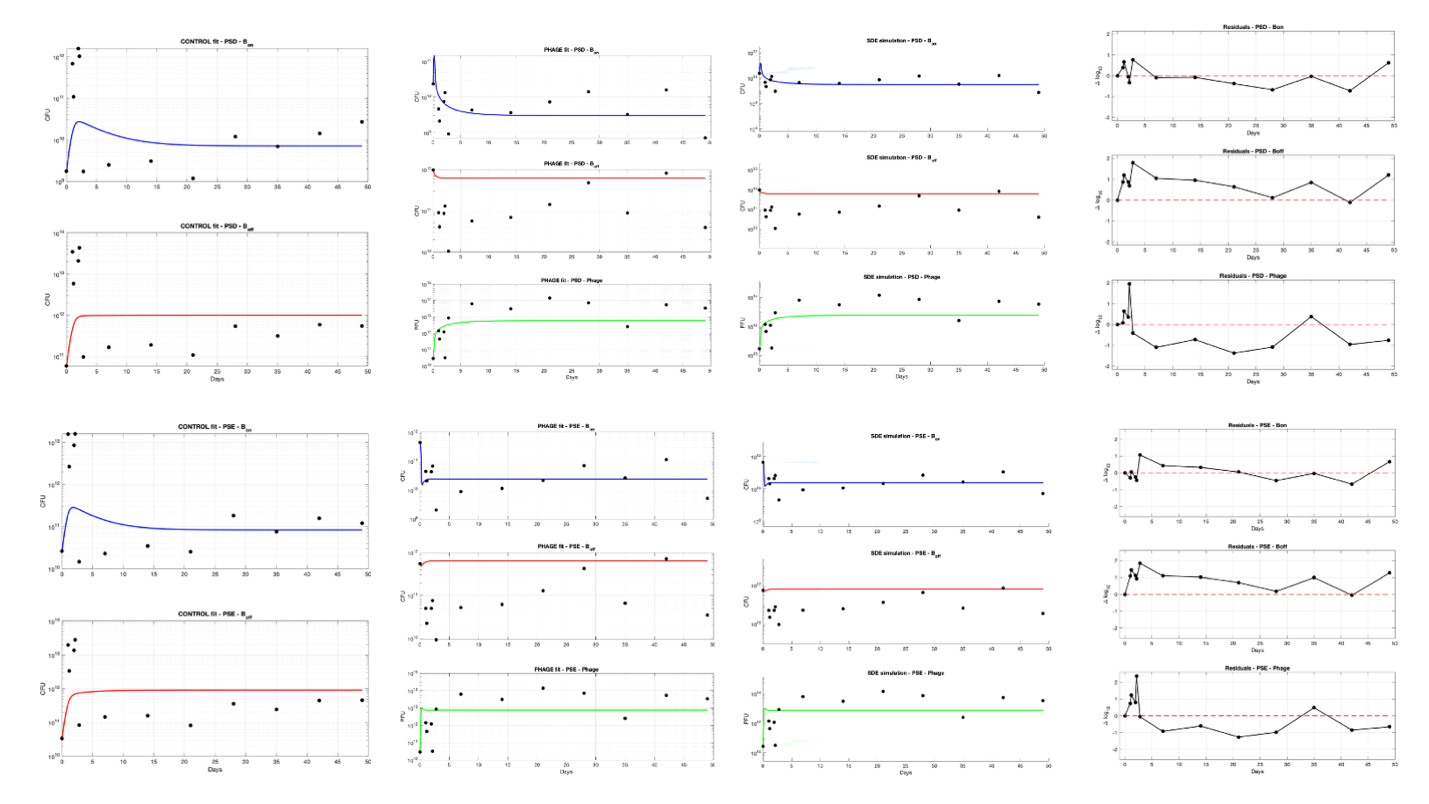

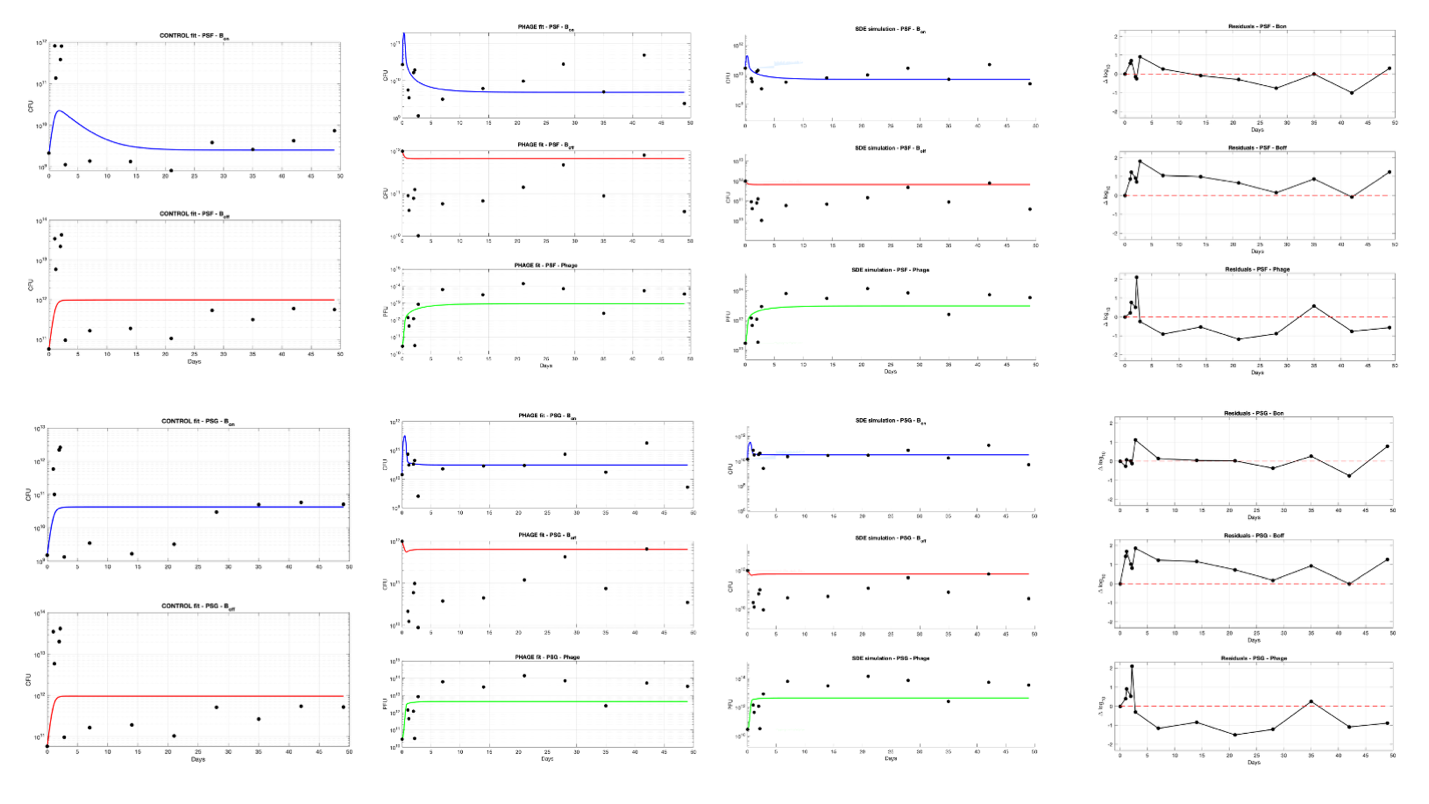

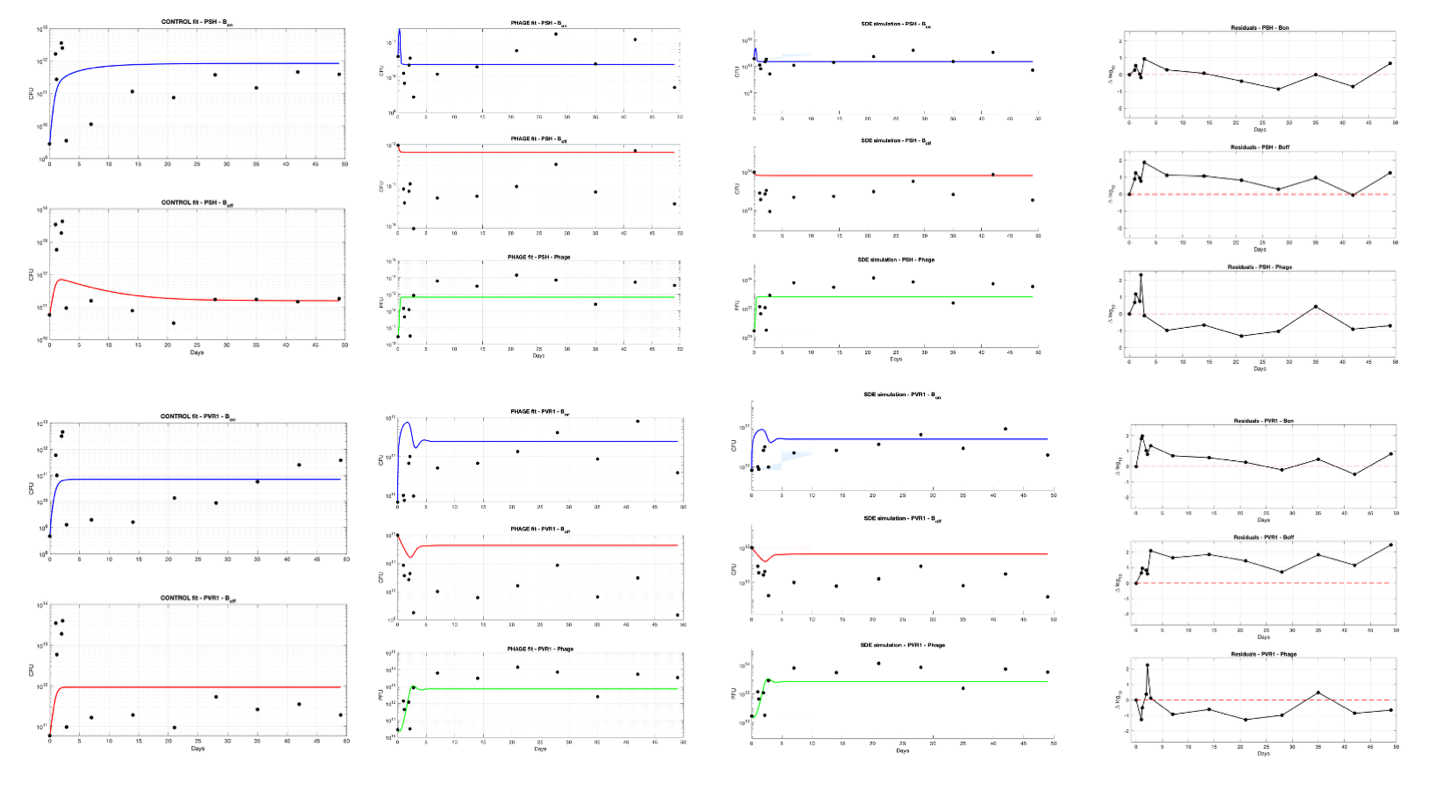

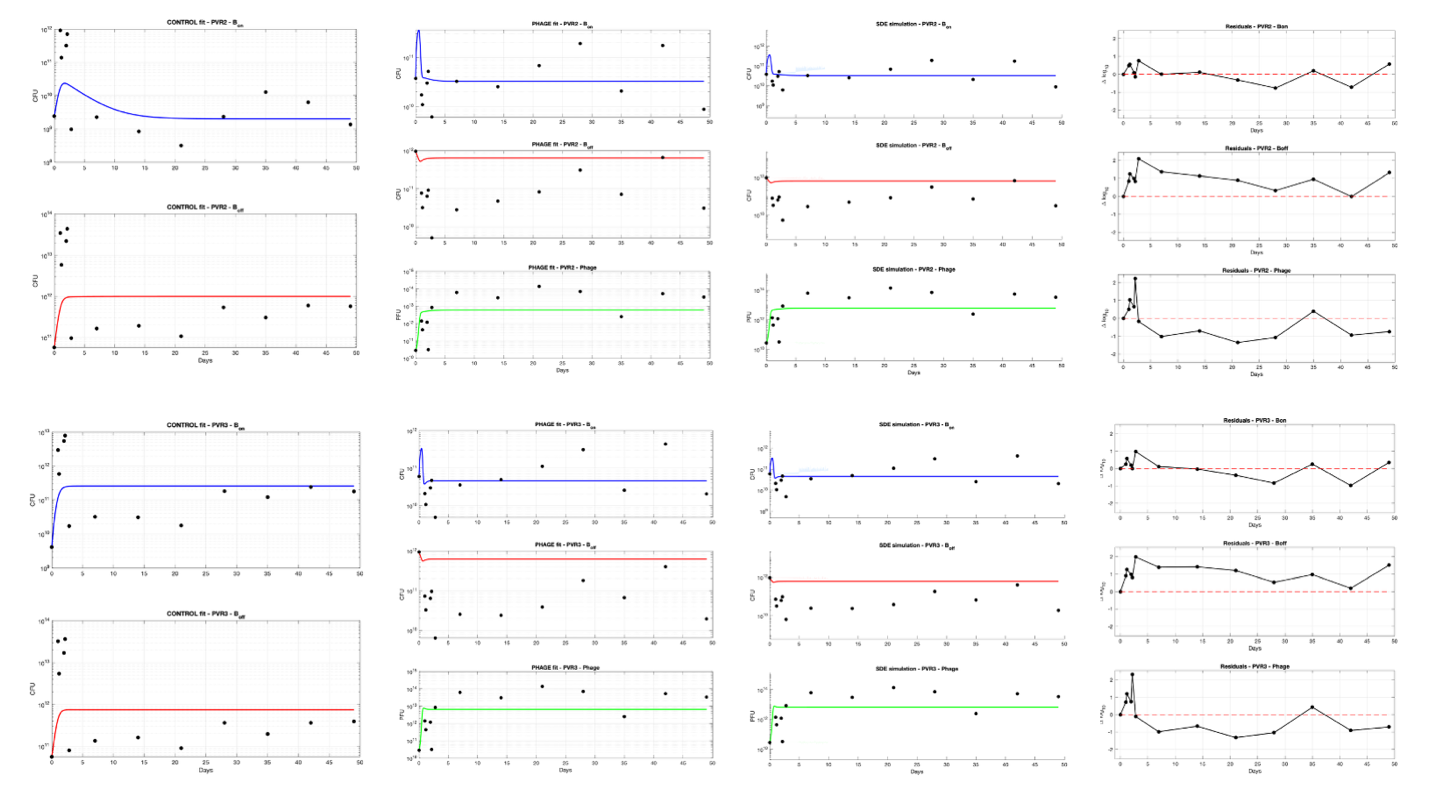

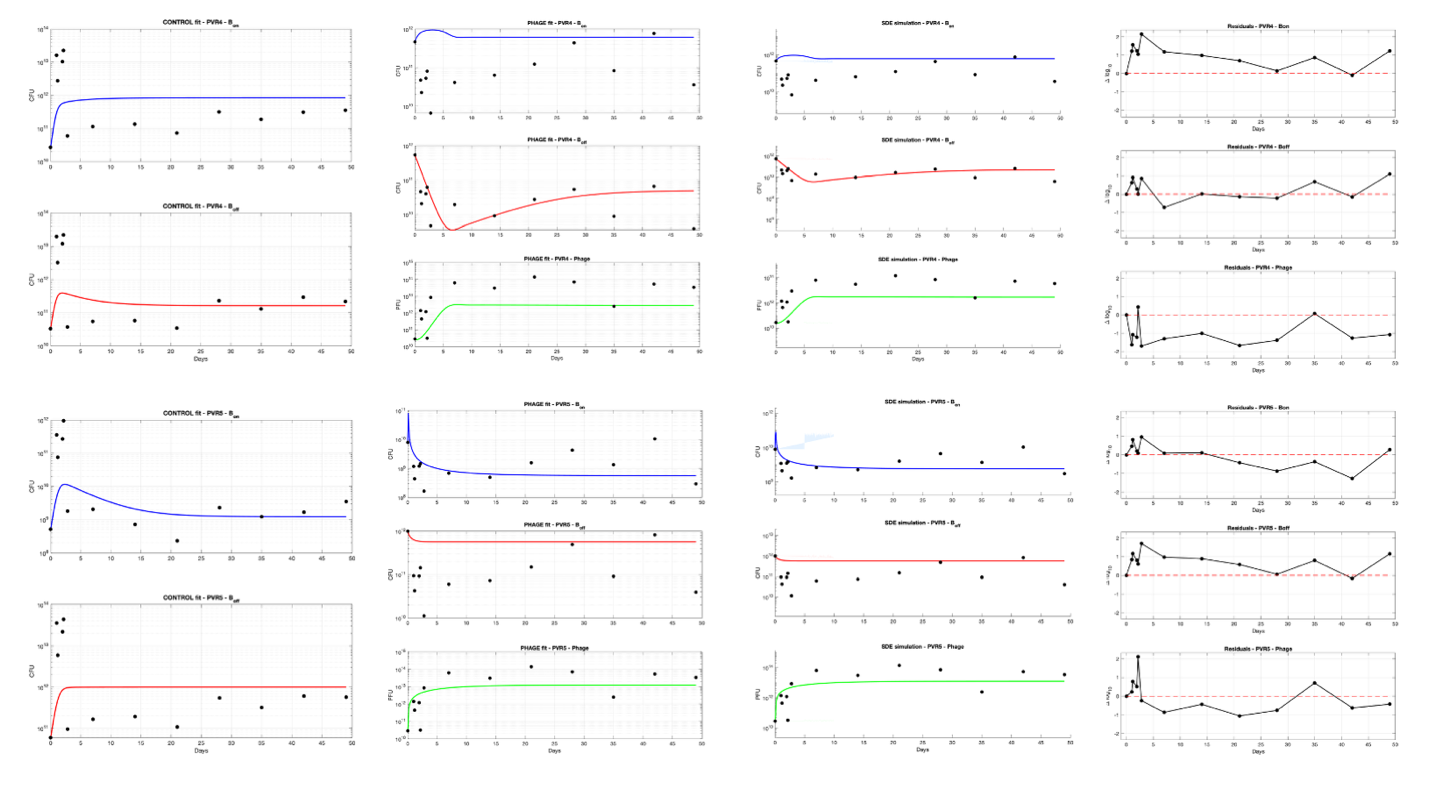

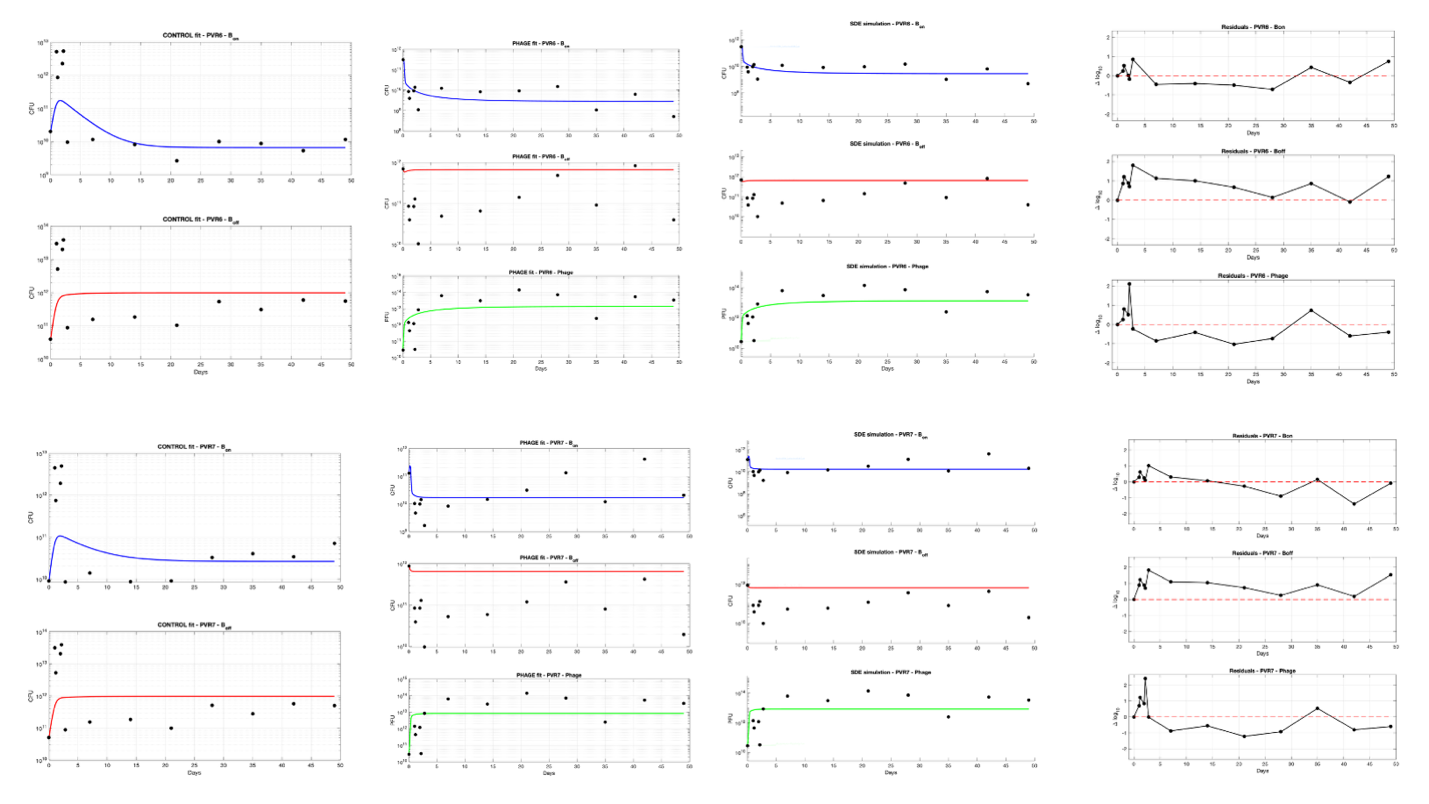

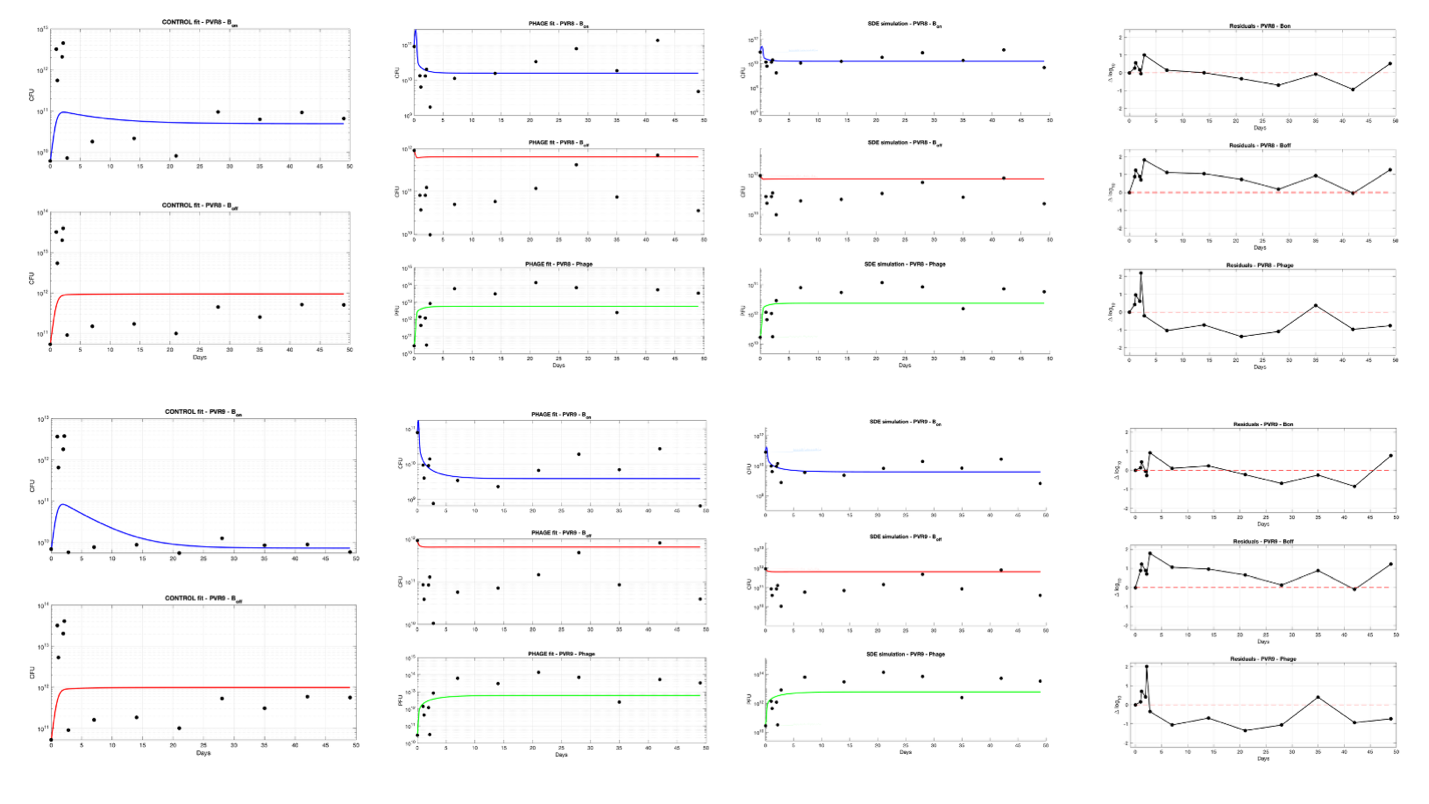

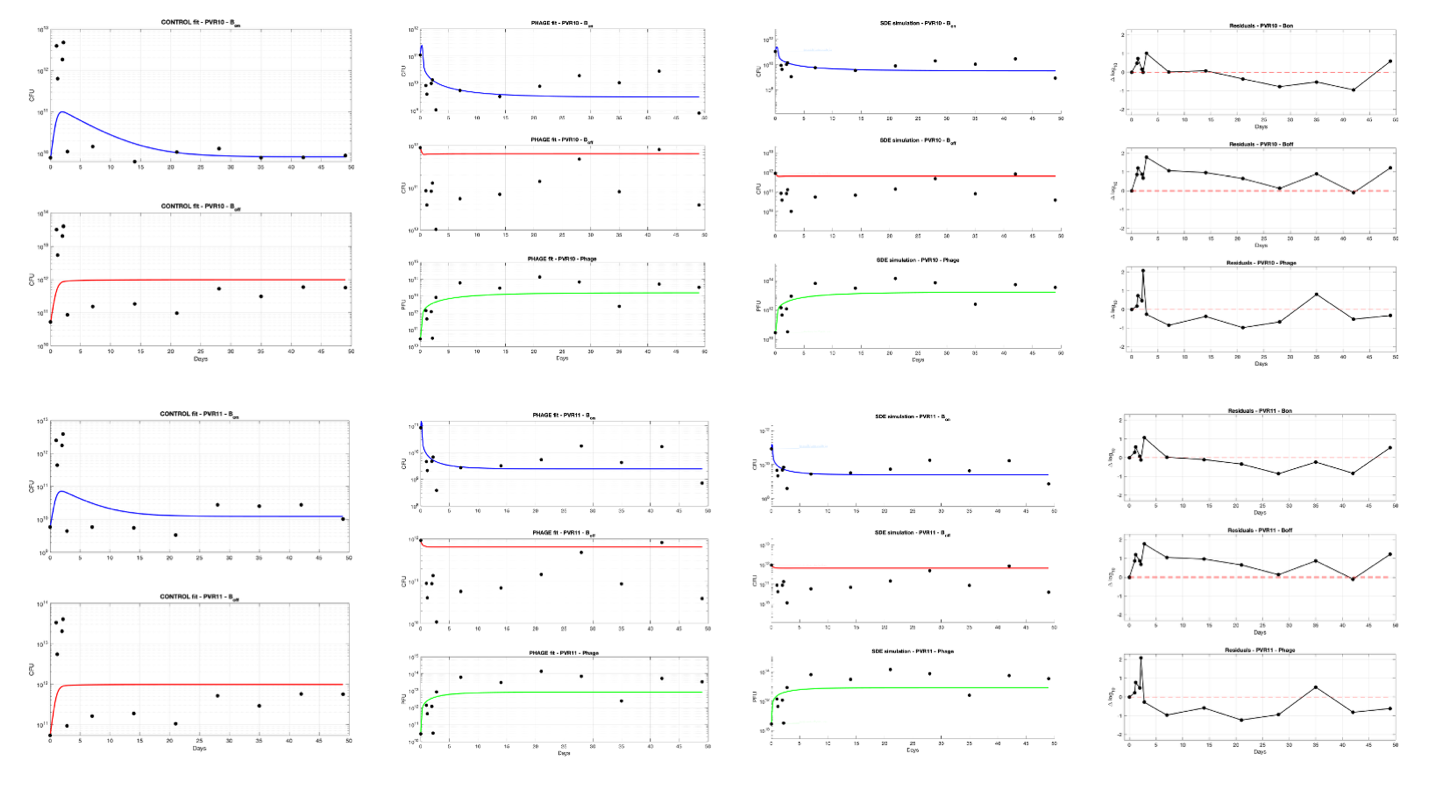

**Supplementary Figure 1. Per-region ODE fits and stochastic validation.**

Deterministic ODE fits, stochastic validation simulations, and residuals are shown for each of the 18 analyzed invertible regions. For each region, control-condition fits are shown for reconstructed ON and OFF bacterial abundances, and phage-condition fits are shown for ON bacteria, OFF bacteria, and phage abundance. Black points indicate observed longitudinal data; colored curves indicate fitted or simulated model trajectories. Residual panels show deviations between observed and fitted trajectories over time, with the dashed line marking zero residual. The figure provides per-locus internal validation of the ODE framework used for downstream region-level susceptibility, contribution-space, and combinatorial analyses.
